# A cerebellar population coding model for sensorimotor learning

**DOI:** 10.1101/2023.07.04.547720

**Authors:** Tianhe Wang, Richard B. Ivry

**Author notes:** Corresponding authors: Tianhe Wang.

## Abstract

The cerebellum is crucial for sensorimotor adaptation, using error information to keep the sensorimotor system well-calibrated. Here we introduce a population-coding model to explain how cerebellar-dependent learning is modulated by contextual variation. The model consists of a two-layer network, designed to capture activity in both the cerebellar cortex and deep cerebellar nuclei. A core feature of the model is that within each layer, the processing units are tuned to both movement direction and the direction of movement error. The model captures a large range of contextual effects including interference from prior learning and the influence of error uncertainty and volatility. While these effects have traditionally been taken to indicate meta learning or context-dependent memory within the adaptation system, our results show that they are emergent properties that arise from the population dynamics within the cerebellum. Our results provide a novel framework to understand how the nervous system responds to variable environments.

## Introduction

Humans are incredibly flexible in how we adapt our motor behavior across variable environments. We readily compensate for the added weight of a heavy winter coat when reaching for an object or adjust the force required as we sip on our morning coffee. The cerebellum is recognized as playing a key role in this adaptation process^1,2^. Impaired adaptation is one of the hallmarks of cerebellar pathology, observed across a range of tasks from experimentally-induced lesions in animal models^3–5^ or neurological disorders in humans^6,7^. Moreover, anatomical and physiological studies have led to computational models in which the cerebellum uses error information to improve subsequent, similar movements^8,9^. This form of learning operates implicitly, automatically recalibrating the sensorimotor system without the need for awareness or drawing on cognitive resources^7,10,11^. The current paper aims to understand the neural computations that support flexible adaptation and how this recalibration process is modified by context and environmental uncertainty.

Previous research has suggested that cerebellum-dependent learning is cognitively impenetrable, responding to error in a “rigid” manner even when the correction fails to improve task performance^10,12–15^. Moreover, unlike many learning processes, adaptation is not sensitive to the statistical properties of the perturbations^16,17^. However, this view of a rigid, inflexible system has been challenged by recent evidence showing that implicit adaptation is modulated by experience^18^. For instance, when participants are exposed to a previously experienced perturbation, the rate of relearning is slower than had been originally observed^18^. Not only does this result suggest a degree of flexibility in adaptation, but, interestingly, this context effect is opposite what is typically observed in studies of relearning: Across a broad range of task domains (e.g., reward-based learning, language acquisition), relearning is typically faster, a phenomenon known as savings^19–21^.

The rigidity and atypical effect of experience point to the need for considering the unique properties of the cerebellum to understand how adaptation is modulated by context and environmental variability. To this end, we develop a novel computational model of the cerebellum. This model incorporates two core observations from cerebellar physiology. First, recent studies of oculomotor control have revealed a fundamental tuning property of Purkinje cells, the primary integrative unit in the cerebellar cortex: These cells are not only tuned to movement direction but also to the direction of error relative to that movement^22–24^. Second, the model includes a two layer network, with the second layer designed to capture activity in the deep cerebellar nuclei^25–28^. We posit that units in the DCN exhibit similar tuning properties as Purkinje cells and that units with similar tuning profiles are linked across these two layers.

We used the model to generate predictions regarding a range of contextual manipulations and evaluated the predictions with a series of behavioral experiments. Specifically, we systematically examined the effect of past experience, error uncertainty, and variation in temporal dynamics in evaluating our model. Where relevant, we consider several alternative models that have been proposed to elucidate how context and environmental uncertainty modulate sensorimotor learning^29–31^. Our population-coding model provides an excellent fit of the behavioral results, even without positing the direct representation of context or latent state variables or having the capability to modulate learning parameters through meta learning. As such, our model provides a comprehensive explanation of core computations that account for how the cerebellum can keep the sensorimotor system well-calibrated across variable environments.

## Results

### Cerebellar Population Coding (CPC) Model

The basic principles of cerebellar-dependent error-based learning are articulated in the classic Marr-Albus model^2,32^. Purkinje cells (PC) in the cerebellar cortex receive two types of input (Fig 1a). One source originates in the pontine nuclei. This pathway is hypothesized to provides contextual information, including an efference copy of the motor command. PCs operate as an internal model, utilizing the input to predict the consequences of the motor command^33,34^. The second source originates in the inferior olive with activation of the climbing fibers indicating a mismatch between the predicted and expected sensory feedback, a teaching signal that is used to update the internal model.

**Fig. 1.**
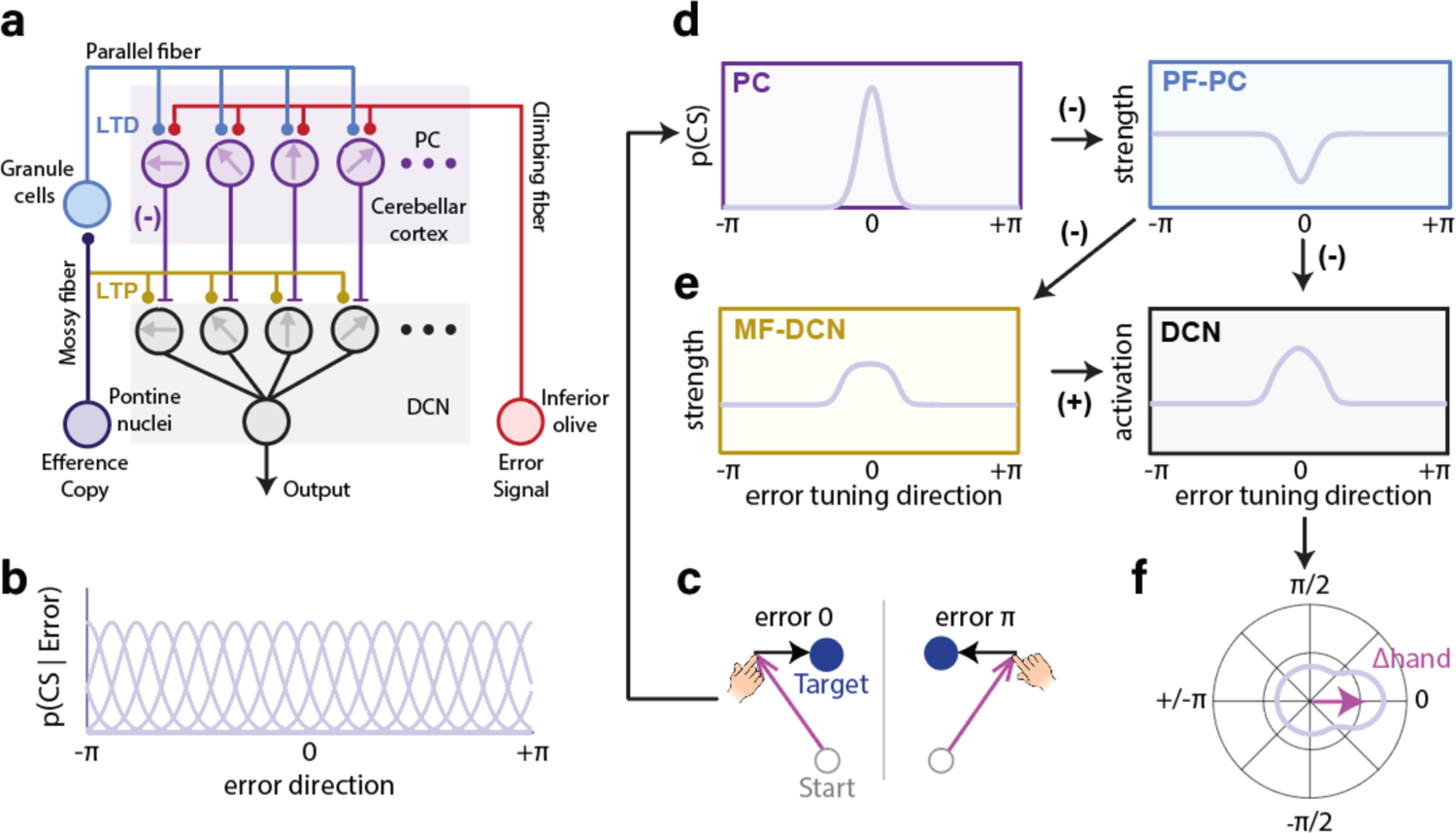
Illustration of the CPC model. **a)** Structure of the cerebellar circuit incorporated in the CPC model. **b)** Each Gaussian-shaped curve represents the tuning function of a single Purkinje cell (PC) based on that cell’s preferred error direction. For the simulations, we used 1000 units with preferred directions that covered 0-π in a uniform manner. **c)** Illustration of visual errors, with the direction of the error specified in polar coordinates. **d-e)** Model-generated adaptation in the cerebellar cortex (d) and deep cerebellar nuclei (DCN) (e). After experiencing an error in 0 direction, PCs with a preferred direction close to 0 will have high probability of generating a complex spike (CS) (d, left) which will result in long-term depression (LTD) for active synapses from granule cell inputs to that PC (d, right). During the preparation of the next movement, the strength of the input from the parallel fibers (PF) will decrease due LTD, attenuating the SS activity of the PC. We assume that attenuation of the inhibitory PC output to the DCN will facilitate long-term potentiation (LTP) resulting from excitatory mossy fiber (MF) input to the DCN (e, left). DCN activation is determined by the excitatory input from the MF and the inhibitory signal from the PC (e, right). The frame colors indicate the corresponding pathway in Panel a. **f)** DCN activation plotted in a polar coordinate. Activation across the population of cells results in a vector (purple arrow) indicating the change in hand angle (Δ hand). Note that the vector points in the same direction as the error (c, left), and thus serves to compensate for the error.

Here we extend the model to provide a general explanation of how learning is modulated by contextual variability. A foundational idea for our model is inspired by a recent work showing how PCs in the oculomotor cerebellar cortex are simultaneously tuned to both movement direction and the direction of a visual error that arises during that movement^22–24^. Tuning in terms of movement direction is reflected in the simple spike activity of the Purkinje cells and tuning in terms of movement error is reflected in the complex spike activity of these cells (Fig 1b). The latter one induces long-term depression (LTD) of parallel fiber-PC (PF-PC) synapses, reducing the future efficacy of similar input on PC activity (Fig 1c-d). Importantly, because the two tuning profiles are in opposite directions^22–24^, error-related activation reduces the simple spike activity, resulting in a change that can reduce the error in the future movement (Fig 1e-f).

A second prominent feature of our model is that plasticity occurs within the cerebellar cortex and deep cerebellar nuclei (DCN)^25,35^. Lesion studies of eyeblink conditioning provide one line of evidence indicating that some aspect of consolidated learning is centered in the DCN. Ablation of the cerebellar cortex can completely block *de novo* cerebellar-dependent learning^26,36^. However, once the learned behavior is established, it can persist after lesions to the cerebellar cortex even though the kinematics are disrupted^37,38^. This dissociation can be explained by the dual-effect of pontine projections to the cerebellum^39^: a polysynaptic projection through the granule layer to PC, and a direct excitatory projection of the mossy fibers to the DCN. We assume that PC and DCN neurons are connected such that they share the same tuning direction for movement^40^, and that learning in the DCN is gated by learning at the cerebellar cortex (Fig 1a). Specifically, LTD at parallel fiber-PC (PF-PC) synapses will reduce inhibitory PC input to the DCN, facilitating the emergence of long term potentiation (LTP) at mossy fiber-DCN synapses (Fig 1e)^41^.

In summary, an error signal will decrease the efficacy of parallel fiber input to PCs and increase the efficacy of mossy fiber input to the DCN (Fig 1d, e). Correspondingly, the net output of the cerebellum will provide a signal of the required change in movement direction to correct for the error (Fig 1f).

### Behavioral Task and Model Parameterization

We first aim to examine the behavior that emerges from the population dynamics of a network in which the individual units are tuned to both movement direction and the direction of movement error. For the work discussed in this section, a single-layer network with this form of representation is sufficient. In the second half of the Results, we will turn to phenomena that motivate a two layered network that maps on to the properties of the Purkinje cells within the cerebellar cortex and the deep cerebellar nuclei.

To measure implicit adaptation, we used a variant of a visuomotor rotation task. Participants reached to a visual target and feedback, when present, was limited to a cursor. To restrict learning to implicit adaptation, we used task-irrelevant clamped feedback in which the radial position of the cursor was locked to the hand, but the angular position was predetermined for each trial. In most experiments, the cursor was shifted by a constant angle relative to the target (Fig 2a-b)^10^. Despite being fully informed of the non-contingent nature of the feedback and explicitly instructed to ignore the feedback, the reach angle gradually shifts in the opposite direction of the clamp^10,18,42,43^. Clamp-induced adaptation has all of the hallmarks of implicit adaptation and, as with other forms of this type of learning, is dependent on the integrity of the cerebellum^10,44^.

**Fig. 2.**
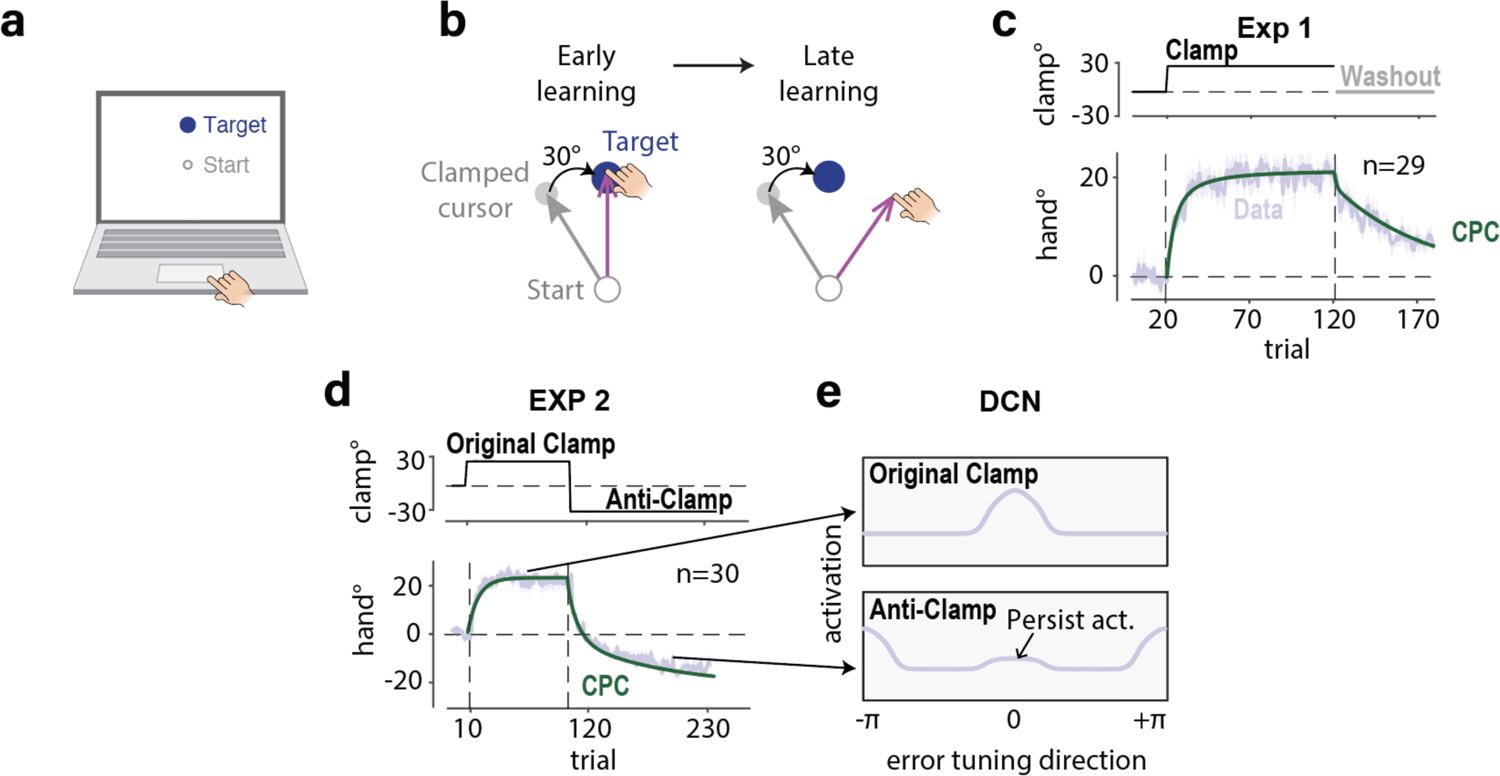
Cerebellar population coding captures learning, forgetting, and anterograde interference during implicit adaptation. **a)** For online testing, stimuli are presented on the participant’s laptop computer and movements are made on the trackpad. **b)** For clamped feedback, the angular position of the cursor is rotated by 30° with respect to the target, regardless of the heading direction of the hand. **c)** Perturbation schedule (top) and results (bottom) for Exp 1. Time course of the mean hand angle is shown in light violet. The CPC model provides a good fit in both the training and no-feedback washout phases. **d)** To examine anterograde interference, the direction of the clamp was reversed during the training section of Exp 2. The behavioral results match the prediction of the CPC, using the parameters estimated from Exp 1. **e)** We mark the direction of a clockwise error (30°) as 0 in the PC/DCN tuning space, and the direction of an opposite (counterclockwise) error (−30°) error as π. Memory of the original perturbation (top row) persists in the anti-clamp training phase, indicated by the activation of neurons tuned to 0 in the bottom row. This residual memory causes anterograde interference. Shaded area in c, d, e indicates standard error.

To determine the learning and forgetting rates for the units of the CPC model, we conducted an experiment (Exp 1) in which participants were exposed to 100 trials with clamped feedback (30°) followed by 60 trials without any feedback. We measured the retention rate from the washout block and then fit the learning rate from the training phase. The data were also used to determine the scaling factor, a parameter that transforms neural activation into hand angle. The CPC model provided an excellent fit for the observed change of hand position (Fig 2c). To rigorously test the model, we fixed the parameters described above when running simulations to generate predictions for the other experiments.

### Anterograde Interference

In the following sections, we will examine several key predictions derived from the CPC model concerning how the experimental context should modulate implicit adaptation. One basic feature of the CPC model is that, for each unit, learning is fast due to the potent impact of complex spikes whereas forgetting in the absence of feedback occurs relatively slowly due to a passive decay process (Fig 2c). Given the slow forgetting, the population activation will be influenced by the persistent activation of units that were tuned to a recent error. As such, when the perturbation direction is abruptly reversed, the model predicts that the observed rate of change in performance will be attenuated due to the persistent activation, even if the learning and forgetting rates are invariant in the model (see Fig 2e).

To examine this model prediction, we used a task in which the sign of the clamp was reversed after an initial training block (e.g., 30° followed by −30°, Exp 2, see Fig 2d). Consistent with the model prediction, the results showed that the rate of adaptation was slower in response to the reversed clamp compared to the original clamp^45–49^. Strikingly, the degree of attenuation closely matched the CPC model’s prediction based on the parameter values estimated from Exp 1. These results indicate the population dynamics within the CPC model can be useful to explain the contextual effects in adaptation.

### Absence of Spontaneous Recovery

Here we aim to compare the CPC model with several alternative models that have been proposed to account for contextual effects in implicit adaptation. Anterograde interference has typically been explained by models positing context-dependent learning mechanisms^30,50,51^. For example, the contextual inference (COIN) model assumes that the motor system forms separate memories for different contexts and chooses which memory to use based on the inferred context^29^. To account for the results of Exp 2, COIN would first build a memory for the 30° perturbation and then a second, distinct memory for the −30° perturbation. Anterograde interference would arise because the introduction of the −30° perturbation would lead to some degree of recall of the response to the initial perturbation. Over time, this would shift to a bias to recall the response to the second memory. In contrast to the COIN model, the CPC model does not posit distinct memories for different perturbations; rather anterograde interference emerges from the population dynamics.

The dual state-space (Dual SS) model^21,39^ offers another account of anterograde interference. This model posits that learning involves two processes that operate at different rates (Fig S1a). When the perturbation is reversed, a fast process will quickly respond to the new perturbation and drive adaption in the opposite direction. However, a slow process will continue to be dominated by its response to the original perturbation, thus producing anterograde interference. In contrast, the CPC model can account for anterograde interference without positing different learning/retention rates across units.

While the COIN, Dual SS, and CPC models make similar predictions about anterograde interference, they make differential predictions on another memory phenomenon, spontaneous recovery. A paradigmatic design to elicit spontaneous recovery in sensorimotor learning studies would be to train participants with a perturbation in one direction, extinguish the adapted behavior by shifting the perturbation in the opposite direction, and then examine behavior in the absence of feedback (Fig 3a)^21^. Spontaneous recovery refers to the fact that the initial movements during the no-feedback phase are in the opposite direction of the initial perturbation (Fig 3b top left). By the COIN model, spontaneous recovery occurs since there is some degree of recall of the original context during the no-feedback phase. By the Dual SS model, the state of the fast process will decay back to zero (i.e., baseline). However, the state of the slow process is still shifted in the direction induced by the initial perturbation, resulting in the manifestation of spontaneous recovery in the no-feedback phase (Fig 3b, S1b). In contrast, the CPC model predicts that spontaneous recovery will not occur when learning is restricted to the implicit system since the model does not have a mechanism for context-dependent memory (Fig 3b bottom right).

**Fig. 3.**
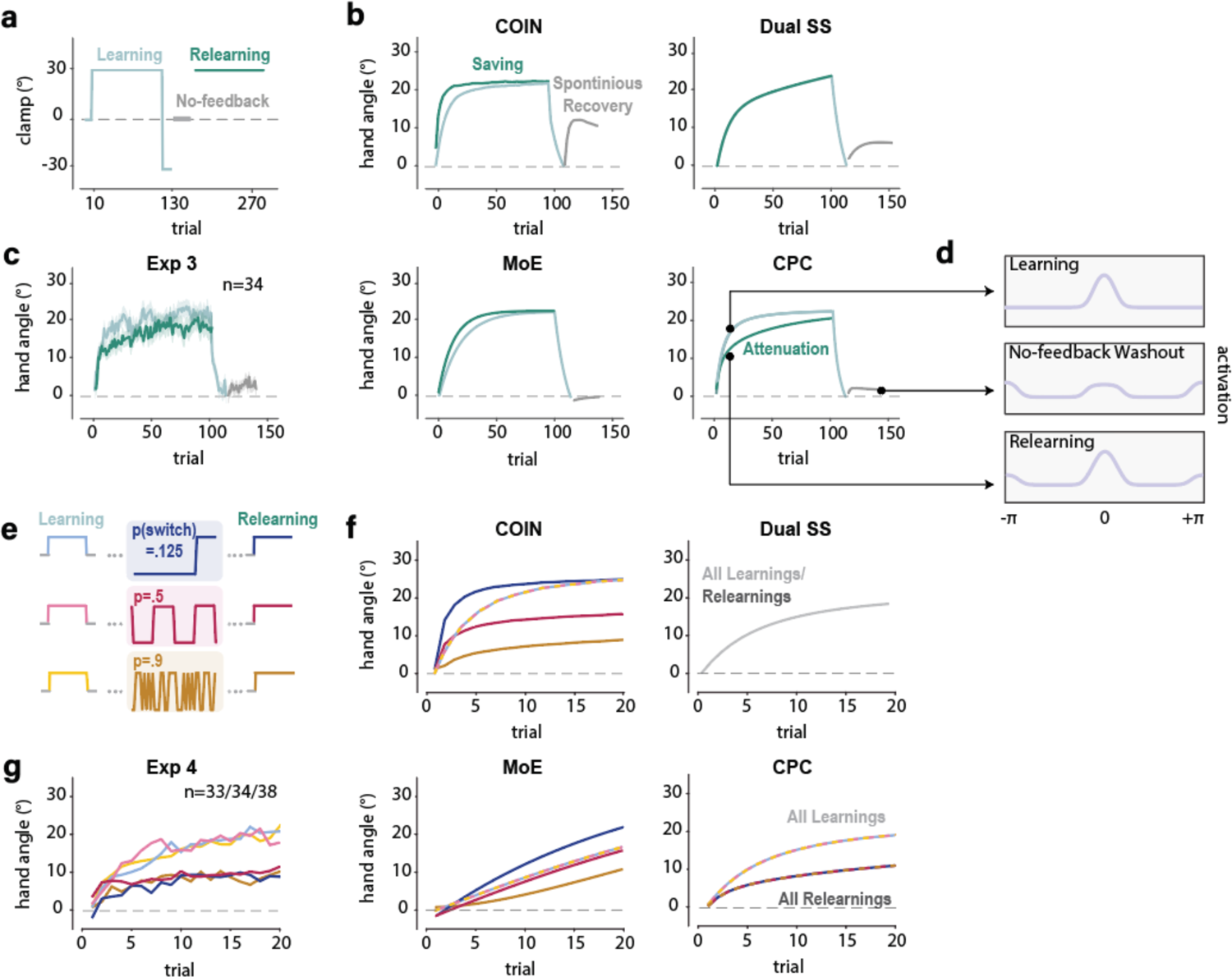
Context as an emergent property of the CPC model. **a)** Exp 3 perturbation schedule. To examine spontaneous recovery, a perturbation in one direction is presented for an extended phase and then reversed for a short phase. The critical test is in the subsequent no-feedback phase. To test for savings, the original perturbation is then reintroduced. **b)** The COIN model predicts spontaneous recovery and savings; the Dual SS model predicts spontaneous recovery but no saving; the MoE model predicts no spontaneous recovery but savings; the CPC model predicts no spontaneous recovery and attenuation upon relearning. Note that, for visualization, the data from the relearning phase are plotted on top of the original learning phase. **c)** Empirical results match both predictions of the CPC model. Shaded area in c indicates standard error. **d)** Population activation in the CPC model at three time points. While the hand angle is close to zero at the end of the no-feedback washout, residual activation resulting from both the original and reversed clamps are evident in the population, indicated by the peak at 0 and π, respectively (middle). The memory of the opposite clamp slowed down the relearning (bottom). **e)** In Exp 4, the learning and relearning phases are separated by a variable phase in which the probability of a perturbation switch is manipulated between participants. **f)** The COIN and MoE models predict that during relearning, the learning rate will be modulated by the prior switching rate (i.e., perturbation variability). The Dual SS model predicts that learning rate will be identical in learning and relearning regardless of the prior switching rate. The CPC model predicts that, while relearning will be slower than original learning, the learning rate will not be modulated by switching rate. **g)** Empirical results are in accord with the CPC model, showing attenuation during relearning and insensitivity to switching rate.

To compare the three models, participants in Exp 3 were trained with a 30° clamp in one direction for 100 trials and then presented with the opposite clamp for 15 trials (Fig 3a). Pilot testing had shown that a reversal of this duration is sufficient to extinguish the shift in hand angle observed to the initial perturbation. The critical test was the subsequent 30-trials no-feedback block. At odds with the prediction of COIN or the Dual SS model, we failed to observe spontaneous recovery (Fig 3c; Fig S2b; t(33)=1.2, p=0.23). While being cautious in drawing inferences from a null result, the absence of spontaneous recovery is consistent with the CPC model.

### Attenuation in Relearning

As probably an even stronger comparison, all three models make unique predictions in the scenario in which the initial perturbation is reintroduced after the no-feedback trials (Fig 3a-b, relearning). The COIN model predicts that relearning should be faster (i.e., exhibit savings) because the system has stored a memory of the initial perturbation. The Dual SS model predicts that the relearning will be identical to the original learning, with no saving or attenuation since the system stores no memory of error. The CPC model predicts that exposure to the opposite error during the washout phase will induce anterograde interference; as such, relearning will now be attenuated. As with spontaneous recovery, the results are at odds with the COIN and Dual SS models, and consistent with the CPC model: Adaptation during the re-exposure block was slower compared to initial learning (Fig 3c; Fig S2a; t(33)=3.1, p=0.004). These results suggest that population activation within the CPC model accounts for contextual effects that cannot be captured by alternative models of sensorimotor adaptation.

### Insensitivity to Error Consistency

The preceding sections demonstrate how the CPC model can account for the dynamics of implicit adaptation in various contexts without positing flexible context-dependent memory like the COIN model. A different form of flexibility concerns the operation of meta-learning processes that modulate model parameters across contexts. This idea is central to the Memory of Error (MoE) model^52^, which posits an optimization the learning parameters based on experienced errors during the training. For example, the learning rate should increase when recently experienced errors are consistent (stable context), and the rate should decrease when recently experienced errors are inconsistent. In contrast, the CPC model does not include a meta-learning process of this sort. Experience-dependent changes in the response to an error will arise because recently experienced errors have transiently altered the population dynamics of the system (e.g., Fig 3d).

To compare the MoE and CPC models, we examined how the consistency of error modulates adaptation. In Exp 4, we tested the response to a clamp with a fixed sign (e.g., 30°) before and after a phase in which the sign of the clamp varied. To manipulate consistency, we varied the switching probability in the variable phase, setting it to 12.5%, 50%, or 90% in a between-subject manipulation (Fig 3e). A key prediction of MoE is that the rate of relearning will be modulated by the switching frequency (Fig 3f bottom). In contrast, the CPC model predicts that the rate of relearning will be independent of switching frequency.

The results were consistent with the CPC model: The learning rate during the relearning phase was not modulated by the switching rate during the preceding variable phase (Fig 3g; Fig S2c; F(2,101)=0.18, p=0.84). Interestingly, relearning was markedly slower during the relearning phase compared to original learning (F(1,101)=37.7, p<0.001). This attenuation is another manifestation of anterograde interference resulting from the opposite errors experienced in the variable-clamp block. We note that these results are not only at odds with the MoE model, but also with the COIN and Dual SS models. As with the MoE model, COIN predicts that the learning rate will be inversely related to perturbation variability (Fig 3f); The Dual SS model predicts that relearning will be identical to that observed during the initial learning phase for all conditions.

To summarize the results from Experiments 1-4, we have examined a variety of tasks used to examine the effect of context on sensorimotor adaptation, comparing the CPC model to three prominent alternative models (Table 1). The results suggest that the population dynamics within the CPC model provide a parsimonious explanation of how perturbation history influences adaption without positing context-dependent memory or meta-learning processes.

**Table 1.** Comparison of the CPC model and Other Models of Sensorimotor Adaptation. Summary of model performance on a set of core phenomena. In evaluating each of the alternative models, we used an implementation based on that presented in the associated paper (recognizing that a reasonable variant might be possible to capture more of the phenomena). The credit assignment model assumes that the agent performs Bayesian inference to decompose the observed error into perturbation sources that vary across different time scales and estimate the optimal policy to compensate for them. ^a^ Both COIN and the MoE models have difficulty explaining why a large change in behavior would be observed in response to random perturbations given their optimality assumptions. One would expect an optimal system to show a minimal or attenuated response to a random perturbation.

|  | CPC | Dual State Space<sup>21,39</sup> | MoE<sup>52</sup> | COIN<sup>29</sup> |
| --- | --- | --- | --- | --- |
| Anterograde interference (EXP 2) | ✓ | ✓ | ✗ | ✓ |
| Attenuation in relearning (Exp 3) | ✓ | ✗ | ✗ | ✗ |
| No spontaneous recovery (Exp 3) | ✓ | ✗ | ✓ | ✗ |
| Invariant to error consistency (Exp 4) | ✓ | ✓ | ✗ | ✗ |
| Slower adaptation after variable perturbation (Exp 4) | ✓ | ✗ | ✗ | ✗ |
| Fast single trial learning for variable perturbation (Exp 4/5/8) | ✓ | ✓ | ✗<sup>a</sup> | ✗<sup>a</sup> |
| Different retention rates for variable and fixed perturbations | ✓ | ✓ | ✗ | ✓ |
| Minimal attenuation of the asymptote in half washout (Exp 6) | ✓ | ✗ | ✗ | ✓ |
| Fast single trial learning around the asymptote (Exp 6) | ✓ | ✓ | ✗ | ✗ |
| Stable process is gated by the volatile Process (Exp 7). | ✓ | ✗ | ✗ | ✗ |

### Tuning properties result in variable changing rates across the population units

A central proposition of the Dual SS model is that adaptation includes two processes with different learning rates (Fig 4a)^21,39,53^. However, within a single layer of the CPC model, an epiphenomenon of population coding is that units will appear to operate in different time scales even if the learning rate parameters are identical across all units (Fig 4b). When an error is observed, cells with a preferred direction centered on this error will display relatively fast learning and quickly saturate (Fig 4c-d). In contrast, cells with a preferred direction slightly misaligned with the error direction will not only learn slower due to the weaker climbing fiber input but will also take longer to saturate. As such, by the CPC model, the change in movement direction is determined by the collective activation of all units, and thus, can be regarded as a composite process of units with different learning trajectories arising from their error tuning.

**Fig. 4.**
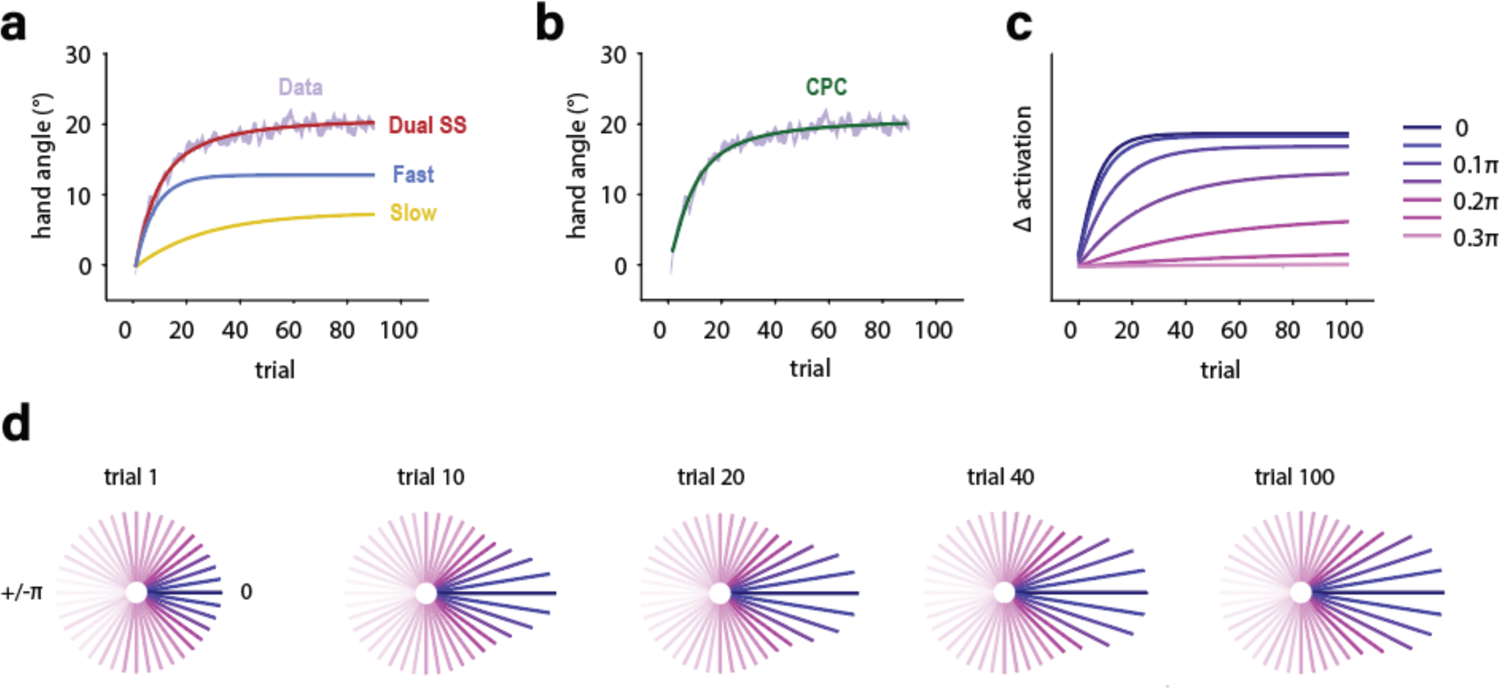
Emergent variation in learning rate across the population units. **a)** Learning curve from Tsay et al (2022) in which participants were exposed to a 30° clamp. This function can be described as sum of a fast process that contributes to the rapid change in hand angle early in learning and a slow process that continues to accumulate over time. The best-fitted Dual SS model and the relative contribution of the fast and slow processes are shown. **b)** CPC model can account for learning function without positing different learning rates. **c)** By the CPC model, the contribution from different units will vary over time due to their tuning for error direction. We mark the direction of the error as 0 in the unit tuning space. Cells tuned to error direction (i.e., 0) respond strongly, driving rapid early learning while saturating quickly. Cells with tuning slightly misaligned with the error direction (e.g., 0.2π) have a small error response but saturate slower; as such they make a relatively large contribution late in training. **d)** Panel c plotted in polar space. Each vector corresponds to a cell with the orientation indicating the cell’s preferred error direction and the length indicating its activation strength.

### Different retention rates in adaptation

We have shown the tuning features of cerebellar units and their population dynamics within the CPC model explain a wide range of contextual effects in implicit adaptation, outperforming alternative models. While we have used a two-layered model in these simulations, the predictions would also hold in a single-layered CPC model. However, the anatomy and physiology of the cerebellum suggest that plasticity effects in the cerebellar cortex and deep cerebellar nuclei can be quite distinct, perhaps constrained by different computational principles^21,39,40^. We explore this issue in the following section, first examining the evidence to suggest computational differences between the layers and then testing predictions to ask how the layers interact with each other.

As shown above, experimental tasks that focus on learning rates are not ideal for assessing the existence and/or dynamics of different layers: A single-layered system can appear to be operating at multiple speeds. As an alternative, we employed a variable perturbation task in which the sign and size of the perturbation were varied across trials (Exp 5, Fig 5a). With this design, the forgetting rate of the system can be empirically measured as the ratio of the change of hand angle in response to the perturbation just experienced (1-back) relative to the previous perturbation (2-back, see Methods). Consistent with previous studies^10,54,55^, we found prominent trial-by-trial adaptation in response to the random perturbations. Surprisingly, the observed retention rate of 0.5 indicates that about half of the learning from the previous trial was forgotten over the 3 s inter-trial interval (Fig 5c). This low value stands in marked contrast with the empirically estimated retention rate from designs in which the perturbation is fixed (e.g., Exp 1, 0.98, Fig 4b,c). When we ran the same analysis on data from published studies using variable or fixed perturbations, we observed a similar marked difference in the retention rate^18,55–57^.

**Fig. 5.**
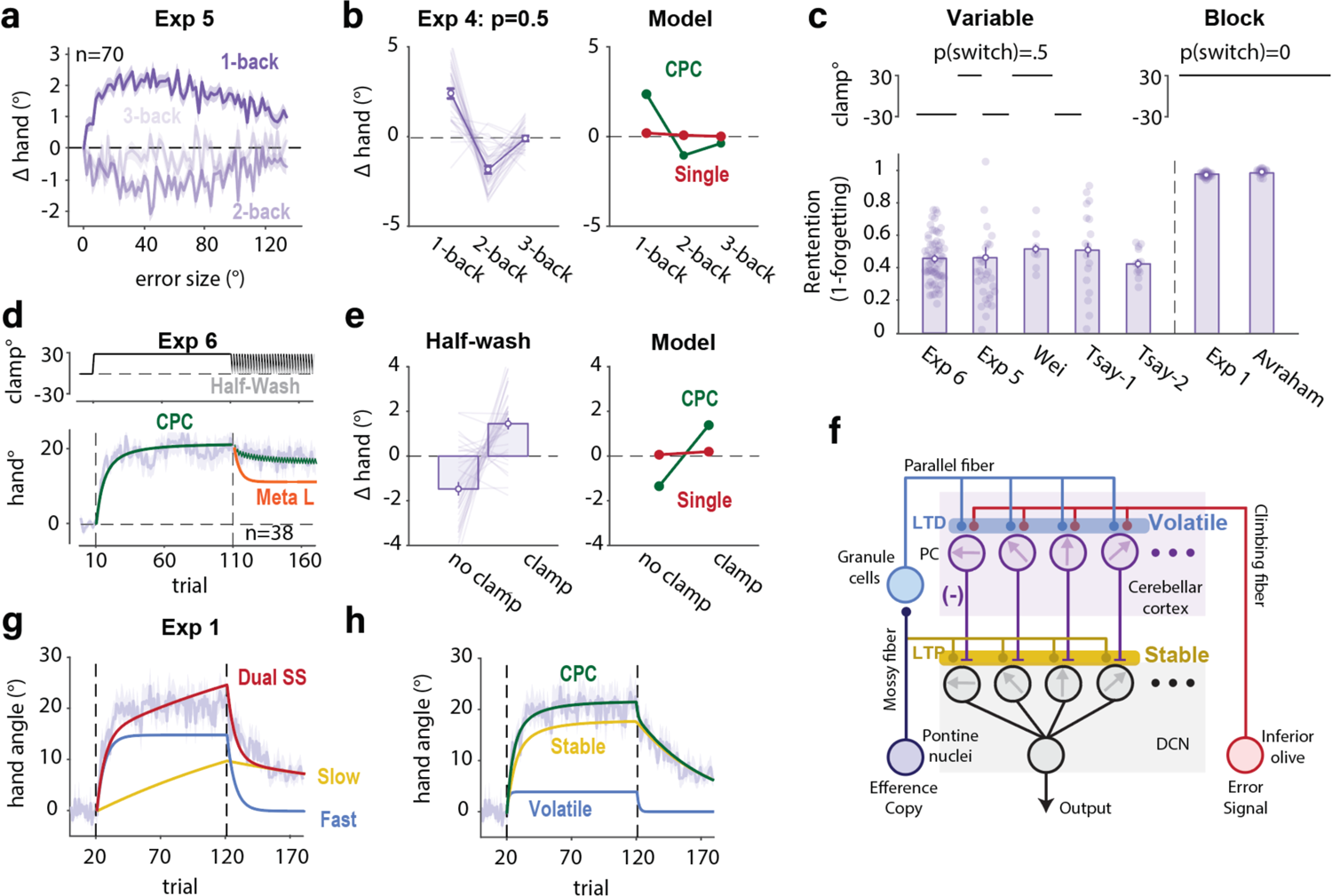
Operation of volatile and stable processes in cerebellum-dependent adaptation. **a)** Trial-by-trial change of hand angle (Δhand) as a function of the perturbation size on trial n-1 (1-back), n-2 (2-back), and n-3 (3-back) in Exp 5. **b)** Left: Similar analysis applied to the data from the variable phase of Exp 4 for the 50% switching condition. Right: The two-layer CPC model can account for the large, yet transient change in hand angle observed in response to a random perturbation. A single-layered CPC model predicts negligible change due to its high retention rate measured from Exp 1. **c)** Estimate of retention rate in experiments using variable or fixed perturbations. Re-analysis of data from Wei: ^55^; Tsay-1: Exp 2 ^56^; Tsay-2: EXP 2 ^58^, Avraham: Exp1 ^43^. All of the depicted experiments used a design with a single target location. **d)** In Exp 6, half washout phase entails a 50/50 mix of clamp and no-feedback trials. Consistent with the CPC model, only a small reduction in hand angle was observed during the half washout phase whereas the meta learning model with a changeable retention rate predicts the hand angle will be reduced by 40%. Purple functions indicate behavioral results. **e)** Large trial-by-trial changes of the hand angle in the half washout phase can only be predicted by the two-layered CPC model rather than a single-layered model. **f)** The volatile process is hypothesized to produce LTD at the parallel fiber-PC synapse; the stable process is hypothesized to produce LTP at the mossy fiber-DCN synapse**. g-h)** The dual rate version of the CPC model (h) provides a better fit of the learning function in Exp 1 compared to the classic dual SS model (g).

This result could suggest that there are two adaptation processes with different retention rates. Alternatively, it might be taken to indicate the operation of a meta-learning process, where the forgetting rate is ramped up in response to a variable environment. To differentiate those two hypotheses, we ran an experiment (Exp 6) in which we first employed a fixed perturbation for an extended block and then followed this with a half-washout phase in which 50% of the trials had no feedback and 50% had clamped feedback (Fig 5d). Interestingly, there was a considerable drop in hand angle after each no-feedback trial, much larger than what would be predicted by a single-layered model parameterized with a retention rate estimated from a typical fixed design (Fig 5e). Indeed, the magnitude of those single-trial changes is comparable to what is observed in a variable design. A meta-learning model would attribute the drop in retention rate to the variable context (namely, the mixture of no-feedback and clamp trials). However, at odds with this hypothesis, the asymptote in the half-washout block largely persisted (Fig 5d). Thus, the retention rate of the system appears to have remained fixed between the learning and half washout blocks. As such, the large trial-by-trial changes in hand angle and a persistent asymptote are consistent with the dual rate hypothesis.

### Differential Plasticity in the Cerebellar Cortex and DCN

Physiological studies have shown that learning in the cerebellar cortex appears to be more stable compared to the DCN. For example, learning-induced changes in the firing rates of simple spikes in Purkinje cells may decrease after a few trials^39,59,60^ whereas changes in the DCN typically last for days^25,35^. Given the evidence reviewed in the previous section pointing to the existence of a system composed of units with distinct retention rates, we instantiated this difference in the two-layer CPC model such that plasticity within the cerebellar cortex/PC can produce changes that are weakly retained whereas plasticity within the DCN is more persistent (Fig 5f and S3).

This conjecture bears similarity to that proposed in previous dual rate models (e.g., Dual SS model)^21,39,61^. However, there are significant differences between these previous models and the two-layer CPC model characterizes the dynamics of the two processes. First, a key feature of state-space models is that adaptation reaches an asymptote when the trial-by-trial effects of learning and forgetting cancel each other out. However, it is difficult to simultaneously fit both the acquisition and washout phases in response to a fixed perturbation with a state-space model, even with the degrees of freedom conferred by a dual rate variant (Fig 5g). In contrast, we posit that learning saturates because of a limit to neuroplasticity within the units^54^. Given this assumption, the CPC model can readily fit the full learning and forgetting function in adaptation (Fig 5h).

A stronger comparison of the two models is provided by the half-washout phase of Exp 6. The SS model predicts that during this phase, the asymptote will eventually drop to 50% of the original asymptote because learning occurs in just 50% of the trials, those with feedback (Fig S6). This prediction holds even in state-space models that posit learning at multiple time scales^21,62^. However, as noted above, the asymptote only showed only a slight decrease when clamped feedback was presented in 50% of the trials, consistent with the predictions of the CPC model (Fig S6b).

A second difference is that, unlike the state space model, the population dynamics within the CPC model enabled it to account for various context effects discussed in Exp 2-4 (See Table 1). Moreover, by extending the dual coding feature in a two-layered system, the CPC model predicts a novel context effect in a random design experiment, one in which the sign of the perturbation is randomized: The trial-by-trial change in hand angle will decrease over the course of an experimental session even if the learning/retention rates of both layers are fixed. Early in the session, learning within both a volatile and stable layer will contribute to the change in performance. However, gradual changes will accumulate in units tuned to both directions within the stable layer, eventually reaching their asymptotic level of plasticity. Thus, late in the session, learning is essentially dependent solely on the rapid changes occurring within the volatile layer. The net effect is that the overall learning rate will decrease over the course of training (Fig S7a-b), another example in which an apparent change in learning rate is emergent from the dynamics of the system. In contrast, the dual state-space model predicts that the overall learning rate should be constant since only the fast process makes a significant response to random perturbations. The data are again consistent with the CPC model: The overall learning rate decreases over the course of a block of trials in which the size and direction of the perturbation is randomized (Fig S7c).

### Learning within the DCN is gated by the cerebellar cortex

Given that the output of the PCs is the primary input to the DCN, we can ask how learning within the DCN is modulated by activity in the cerebellar cortex. By the CPC model, learning in the DCN is scaled by the change in simple spike activity of the PCs; in effect, the stable process is gated by the labile process. While this hierarchical organization has been proposed previously in the discussion of dual-rate models of adaptation^39^, it has not been tested empirically.

One way to test the gating hypothesis is to manipulate the duration of the inter-trial interval. The change in hand angle arising from a volatile process decreases with the passage of time. If the volatile process gates the stable process, increasing the ITI should result in a much slower change in hand angle in a fixed design (Fig 6b). In contrast, if the two processes operate in parallel (PARALLEL model), the operation of the stable process will not be influenced by variation in ITI. Higher asymptotic values in a short ITI condition would be solely due to the greater contribution of the volatile process.

**Fig. 6.**
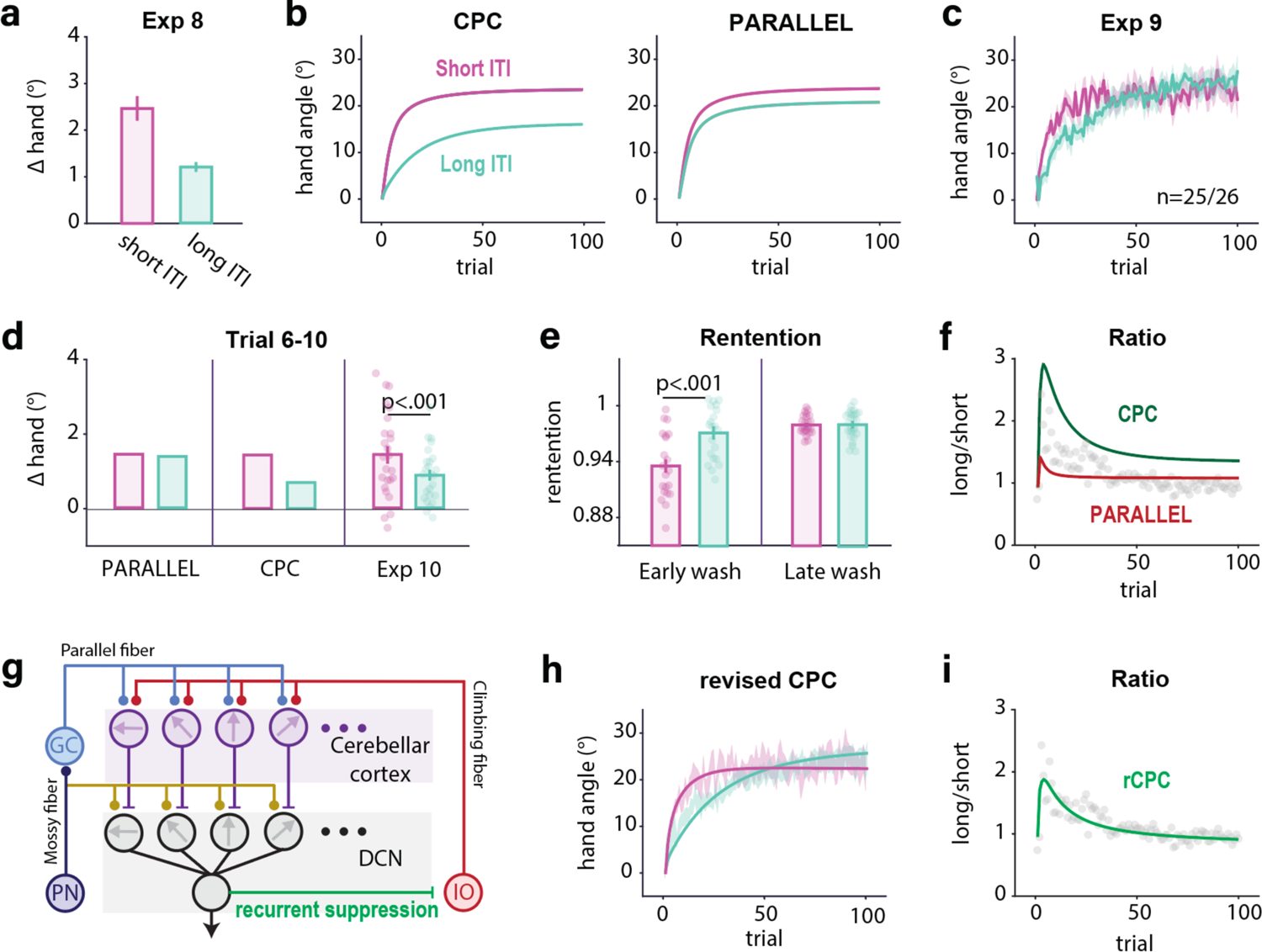
The stable process is gated by the volatile process. **a)** Trial-by-trial change in response to a variable perturbation with a short (data taken from Exp 4, p(switch)=0.5 condition) or long ITI (Exp 8). **b)** Predictions of learning functions under the gating assumption of the CPC model and alternative model in which the two processes operate independently (PARALLEL). **c)** Learning functions in Exp 9 using either a short or long ITI. Consistent with the CPC model, the difference between the two functions is reduced over time. **d)** Model predictions and results from Exp 9 for the change of hand angle across trials 5-10. The change of hand angle is higher in the short ITI condition. **e)** The retention rate is larger in the initial no-feedback trials in the long ITI condition since the volatile process is weakened by the passage of time. However, the retention rate is similar across the two ITI conditions late in washout, consistent with the hypothesis that only the stable process remains operative. **f)** Hand angle ratio between short ITI and long ITI conditions deviates from the predictions of both models. The ratio falls between the two model predictions early in training and is smaller than predicted by both models late in training. **g)** Revised CPC model includes inhibitory projection from DCN to the IO. This suppresses the error signal conveyed by the climbing fibers. This suppression is assumed to decay with time, becoming negligible in the long ITI condition in the revised CPC model. **h-i)** Predictions of the revised CPC model provide a good fit to the learning curve (h) as well as the change in the ratio between the long and short ITI conditions (i) in Exp 9. Shaded area and error bar indicate standard error.

To evaluate the gating hypothesis, we employed a variable design in Exp 8, using a 7 s ITI. By comparing this long ITI condition to a short one (Exp 4) which involved the same design but in which the next trial started as soon as the hand was repositioned at the start location (0 s ITI), we estimated the decay rate of the volatile process across time. Based on the estimated parameter, we predicted the learning function in response to a fixed perturbation (Exp 9), again comparing a 7 s and 0 s ITI conditions (Fig 6b). To measure learning within the stable process (DCN), we ignored the first five trials since these would have a significant contribution from the volatile process. Over the next five trials, we found that the change in hand angle was much higher in the short ITI condition (Fig 6d), suggesting that the learning in the stable layer (DCN) is modulated by the volatile layer (PC).

Furthermore, consistent with the prediction of the CPC model, the long ITI condition showed a slower decrease in hand angle compared to the short ITI condition in the initial washout trials (Fig 6e), suggesting a smaller contribution of the volatile process in the long ITI condition. This difference diminished in late washout trials since the residual memory here comes from the state of the stable process.

However, we note that this version of the CPC model fails to capture one prominent feature in these data, the convergence of the two functions at asymptote (Fig 6c & f). The CPC model predicts that the advantage for the short ITI condition should persist, resulting in a lower asymptote in the long ITI condition. We modified the CPC model (Fig 6g), adding an inhibitory connection from the DCN to the inferior olive, a pathway that has been identified in physiological studies^63,64^. The inhibitory signal to IO will reduce its input strength to the cerebellar cortex, thus decreasing the number of PCs that generate complex spikes^65^. Assuming the strength of this inhibition decays across time (ITIs), more learning units can be recruited in the long ITI condition though plasticity within each unit is reduced due to the decay of memory during the long ITI (Fig S5). The revised CPC generates learning functions that provide good fits in both ITI conditions (Fig 6h & i). Importantly, after re-estimating all of the parameters using this variant of the CPC, we observed negligible effects on the predictions reported for the other experiments (Fig S8). In sum, these results point to a hierarchical arrangement in which the volatile process gates the operation of the stable process.

The revised two-layer CPC model can account for another classic learning effect, contextual interference, referring to the phenomenon in which performance gains are slower when training involves multiple contexts (e.g., reaching towards multiple directions) compared to training in a single context, but retention is better in the former^69,70^. This phenomenon, at least in the context of implicit adaptation, is an emergent property of the parallel operation of volatile and stable learning processes (see Supplementary result 1, Fig S9).

## Discussion

To support flexible behavior, an organism needs to choose an action appropriate for a given context and execute a movement to achieve the desired outcome. A large body of work has sought to delineate the principles of these learning processes, with one prominent question centering on how the processes incorporate context and respond to uncertainty. Here we address this question with respect to the cerebellum, a subcortical structure long recognized as essential for keeping the sensorimotor system precisely calibrated in the face of fluctuations in the environment or state of the agent. We developed a population-coding model incorporating two key features: 1) Units are tuned to both movement direction and error direction, and 2) learning occurs at different rates in the cerebellar cortex and deep cerebellar nuclei, with the former characterized by a fast, volatile process and the latter characterized by a slower, stable process. Our CPC model provides a parsimonious account of a diverse range of learning phenomena and offers new insight into the temporal dynamics of learning (Table S1).

### Context Dependency as an Emergent Property of Population Coding

Importantly, there is no explicit role of context in the CPC model in the sense that a context does not trigger the retrieval of its associated response. Rather the signatures of context-dependent learning and environmental uncertainty emerge naturally from a population of tuned elements that operate in an inflexible manner. In this way, the CPC model diverges from classic models in the behavior that emerges when a previously encountered context is re-experienced. Under such circumstances, classic models predict savings in relearning given that the context facilitates the retrieval of the appropriate response^29^. In contrast, the CPC model accounts for the fact that when a previously experienced perturbation is encountered, implicit adaptation not only fails to show savings, but actually can show attenuation^18^. This attenuation can be seen as another manifestation of anterograde interference: Due to the tuning properties of neurons in the cerebellar cortex and DCN, persistent activation in response to error in one direction will interfere with the response to an error in a different direction.

Given the impressive flexibility in human motor learning, it might be surprising that implicit adaptation does not explicitly track the context or uncertainty of the environment^29,50,66^. We propose that this rigidity reflects a degree of modularity between processes associated with action selection and those related to movement implementation. The cerebellum is part of a system designed to use error information to ensure the accurate execution of a planned movement. The emphasis here is on “planned movement” rather than “desired outcome” to underscore the point that this system appears to operate independent of the task goal; indeed, participants will adapt to sensory prediction errors even when the change in behavior is detrimental to task success^10,12^. This modularity provides a means to keep the system properly calibrated across changes in the internal state of the organism (e.g., perceptual biases, fatigue), factors that need not require a change in action planning. In contrast, other learning systems are designed to use error information related to task success to determine if the selected action was optimal given the current context. These systems would be optimized to track contextual shifts in determining the appropriate policy. Consistent with this hypothesis, contextual effects such as savings and sensitivity to uncertainty are observed in adaptation tasks that benefit from changes in action selection^52,67,68^.

### Hierarchical Organization Within the Cerebellum for Implicit Adaptation

The behavior observed in adaptation studies is assumed to reflect the function of learning processes that operate at different time scales^21,61^. It has been suggested that fast and slow processes correspond to explicit and implicit learning processes^62^. By using variable and fixed designs, we provide evidence that learning limited to just the implicit system operates at different timescales, a notion similar to the original framing of the Dual SS model.^21,39^ However, rather than view these as processes that operate in parallel, our empirical and modeling results highlight a hierarchical organization in which accumulated learning from a volatile process will constrain the learning rate of a stable process. This organization readily maps onto a two-layered network formed by the cerebellar cortex and DCN, with the output from the former gating learning within the latter. The neurophysiological evidence is consistent with this assumption. While the change of SS activation in the PCs can happen within a few trials^22,59^, changes within the DCN can maintain learning across days^25,35^. Reflective of the hierarchical organization, we showed that there is an asymmetric dependency such that the synaptic strength in the cerebellar cortex determines the PC output that modulates learning within the DCN.

We recognize that a two-layered model is clearly a simplification. Indeed, to explain the asymptotic convergence in the long and short ITI conditions, we had to incorporate a third layer into the model, creating a closed loop by adding a projection from the DCN to the IO. While the anatomy supports the existence of this pathway, to achieve convergence, we added two specific features to the dynamics of this pathway. First, the intensity of the inhibition from the DCN to the IO exhibits intensity decrease over time^71,72^. Second, the projection is generic, inhibiting IO units independent of the directional tuning of the DCN neuron. These two assumptions need to be tested in future physiological studies.

### Generalization of the CPC model

Though our model focuses on the cerebellum and sensorimotor learning, the core computational principles may offer insight into how the nervous system responds to environmental uncertainty. The population-coding aspect of the CPC model is similar to models of perceptual learning that include a basis set of tuned elements^73^. For example, in models of time perception, contextual effects on perceived duration have been proposed to reflect the interaction of units tuned to different durations^74,75^. Applied to this domain, the CPC model could be used to derive specific predictions of how temporal perception is modulated by uncertainty and establish boundary conditions for interference. Moreover, the two-layer network in the CPC model provide a novel framework to understand of how learning can involve multiple processes that follow different temporal dynamics, a phenomenon widely observed in cognitive tasks. For instance, value learning has been hypothesize to reflect the joint operation of a fast, volatile process and a slow, stable process^70^. While these processes are typically viewed as operating in parallel, the CPC model offers an example of how a hierarchical framework might prove more parsimonious.

## Methods

### Cerebellar Population Coding (CPC) model

Here we extend the classic Marr-Albus model, focusing on how learning is modulated when the environment is variable. A foundational idea for our model is inspired by a recent work showing how PCs in the oculomotor cerebellar cortex are simultaneously tuned to both movement direction and the error that is associated with that movement ^22,23^.

To examine the implications of these tuning properties for cerebellar-dependent learning, we incorporate PC tuning into a learning model. Specifically, we formalize the teaching signal, the complex spike (CS) activity of a PC with a preferred direction of *i* (0≤ *i* < π) in response to a movement error *e* (Fig 1d) as:

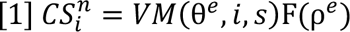

where *VM*(*i*, *s*) is the probability density function of a simplified circular (von Mises) distribution with a mean of *i* and standard deviation of *s*. θ^#^ and ρ^#^ refer to the direction and the size of *e*, respectively, and *n* is the trial number. Given we only applied one error size across all experiments, F(ρ^#^) is not relevant and set as 1 here^55,57^. The form of this non-linear relationship is not relevant in the experiments with a fixed perturbation; for those experiments, the exponent was set to 1.

Following the Marr-Albus model, the occurrence of a CS suppresses the strength of the parallel fiber input synapse (*W*) through long-term depression (LTD):

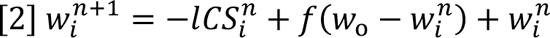

where *l* (*l* > 0) and *f* (0 < *f* < 1) are the learning and forgetting rates, respectively, and *W*_&_ is the baseline synaptic strength. Since the level of single spike (SS) activity will be greatest for cells coding a movement direction opposite to the error, the modulation of synaptic strength will drive the next movement in a direction that corrects for the observed error.

The preceding paragraph describes how parallel fiber synapses onto PCs are modified. A second prominent site of plasticity is at deep cerebellar nuclei^25,35^. Importantly, PC and DCN neurons are organized such that they share the same tuning direction for movement^40^. We posit that learning at the DCN is gated by learning at the cerebellar cortex. Specifically, LTD at parallel fiber-PC (PF-PC) synapses will reduce inhibitory PC input to the DCN, resulting in long term potentiation (LTP) at the mossy fiber-DCN synapses (*m*) (Fig 1e):

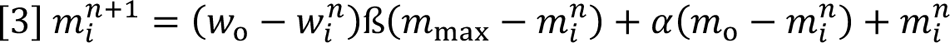

where ß and α are the learning rate and the forgetting rate of the DCN input synapse, respectively. The parameters *m*_&_and *m*_’()_ represent baseline and maximal synaptic strength, respectively. The latter constraint is based on empirical results showing that implicit adaptation saturates independent of the error size.

In this initial version of the CPC model, we have assumed that the PC layer is dominated by LTD, and the DCN layer is dominated by LTP. Classically, PC-layer learning has emphasized LTD^1,2,76,77^, including recent evidence from rodent work showing LTD during upper limb reach adaptation^3^. There is a dearth of evidence concerning the mechanisms of learning in the DCN. As such, we assumed that learning here follows a simple Hebbian process, one consistent with LTP. Importantly, our assumptions regarding LTD and LTP are not critical computationally. The results would be similar if we assumed the reverse or had a mixture of LTD and LTP at each layer. Indeed, a more realistic model should incorporate some degree of bidirectionality given that LTP is also observed in the PC layer^23,60^. For example, an error signal at 0 could result in units with a preferred direction at π becoming strengthened by LTP^22^, presumably because of their co-activation with a shared parallel fiber input. We expect a model with bi-directional modulation would have greater flexibility. However, for the present purposes, the predictions would be largely unchanged and thus, we opted to go with a simple dichotomy.

Considering the two sites of plasticity, DCN activity on a repeated trial following a movement error can be formalized as:

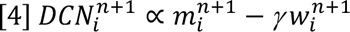

where *γ* is a scaling factor. The output of the population of DCN neurons will correspond to the change in movement direction in response to an error, a signal that can be used to adjust the movement. This can be expressed as: (Fig 1f):

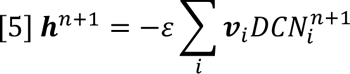

where *h*^n^ is a vector representing the hand angle on trial n, *v*_i_ is a vector representing the tuning direction of unit *i*, and ε is a scaling factor to transfer the neural activity into hand angle.

We note that we do not specify, for the purposes of this paper, whether the cerebellum is best viewed as a forward or inverse model. The current implementation of the CPC model most resembles an inverse model given that the system modifies the motor commends based on the error. However, the model could be reframed as a forward model with the output a prediction that is fed into an (cerebellar or extracerebellar) inverse model that refines the motor commands

### Parameterization of the CPC Model

In our simulations of PC and DCN neurons, we modeled 1000 units for each layer and set the standard deviation of the tuning function (*s*) to 0.2π. The results of most simulations were not sensitive to these two parameters. While anatomical studies show considerable convergence from the PC layer to the DCN, for simplicity, we opted to impose a one-to-one connection between the PC and DCN.

We used an empirical approach to estimate the learning and forgetting rate for PF-PC synapses, using the data from Exp 4 in which +/- 30° clamps were presented with a 50% switching probability. To measure single trial learning, we calculate the change of hand angle between trial n and trial n-1, flipping the sign when the clamp on trial n-1 was negative. To measure single trial forgetting, we calculate the change of hand angle between trial n and trial n-1, flipping the sign when the clamp on trial n-2 was negative.

PF-PC forgetting (*f*) is the ratio of single-trial forgetting and single-trial-learning. By definition, retention rate is 1-(*f*). We applied the same method to measure the retention for all variable designs and this gave us an *f* around 0.5. Model simulations indicate that this method can precisely estimate retention when the perturbation is random. In all of the analyses, we excluded the first 50 trials since learning at this early stage is influenced by both PC and DCN. For comparing the learning rate between early and late training in a by variable design, we employed the same general approach but limited the analysis to the first 50 trials to estimate early learning (Fig S8).

The baseline and maximal strength of MF-DCN synapses can be set to arbitrary values: We used 1 and 1.85 for *m*_&_ and *m*_’()_, respectively. We measured the retention rate of the MF-DCN synapse (α) empirically using the data from the no-feedback washout phase in Exp 1:

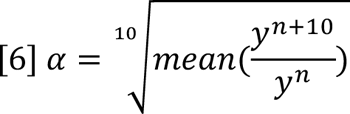

where *y*^n^ is the hand angle in trial n. The first 20 trials in the washout phase were excluded since they may be contaminated by a volatile process.

The learning rate of the PC (*l*) and DCN (ß) and the scaling factors (*γ*, ε) were jointly fitted from the learning block in Exp 1 and the single trial learning in Exp 4. This results in a set of parameters: *l* = .05, *f* = .018, ß = 2, α = .5, *γ* = 0.15, ε = 130. These parameters were fixed in the simulations of all the other experiments. The only exception is Exp 9, where we set the PF-PC retention rate for the long ITI conditions (*f*^+^) to be 0.3, based on the empirically observed value in the variable design of Exp 8.

### Revised CPC model

The results of Exp 8 led us to develop a post-hoc variant in which the output of the cerebellum modulates the input, an idea that is consistent with cerebellar anatomy and physiology^63,78^. The basic version of the CPC model predicts that learning in a long ITI condition will reach a lower asymptote compared to a short ITI condition. This occurs because the contribution of the volatile process is suppressed in the long ITI condition. However, the results of Exp 9 showed that, with a sufficient number of trials, learning in the long ITI condition eventually reaches the same asymptote as in the short ITI condition. This observation led us to revise the model by adding an inhibitory pathway from the DCN to the inferior olive^63,78^.

We assume that the output of the DCN integrates the activation of directionally tuned units and that this signal serves as a generic inhibitory signal to the inferior olive. We implemented this generic suppression by subtracting a common value from the activation of cells tuned to all error directions in the inferior olive (IO):

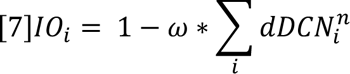

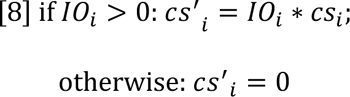

where ω represents the strength of suppression. Given the assumption that ω decreases across time, we used separate parameter values of ω for the long and short ITI conditions ∑_i_ *dDCN*^n^_i_ is the sum of the change of all NCD units relative to their baseline activities. *Cs*’_i_ is the corrected CS activation value after taking the DCN-IO pathway into the consideration and replaces the *Cs*_i_ term in EQ [1-5]. The retention rates of the volatile and stable processes (*f*, α) in the revised CPC model were set as in the basic two-layer model. The other parameters (*l*, ß, ε, ω) were jointly fitted from two data sets, the learning block in Exp 1 and the variable condition in Exp 8. The parameter set is as follow: *l* = .1, *f* = .018, ß = 2, α = .5, *γ* = .2, ε = 210, ω(*short*) = 2.5, ω(*long*) = 0.

### Alternative Models for Comparison

#### Variants of the CPC Model

To help clarify the importance of a two-layer model, we describe two variants of the CPC model. First, we implemented a single-layer version of the CPC model by modifying Eq 4 to:

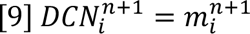

In this version, the output of the system is solely determined by the strength of the MF-DCN.

Second, we implemented a model in which the volatile and the stable processes operate in parallel (PARALLEL) rather than hierarchical as in the CPC model. Since the stable process is insensitive to ITI, we estimate the MF-DCN synapse (*m*) by simulations using a short ITI. The simulated value was then used in simulations of the long ITI condition. For the volatile process, the strength of the PF-PC synapse (*W*) was measured separately for the two ITI conditions.

#### State-space model

We employed a standard version of a state-space model^21,79^:

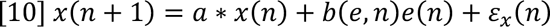

where *x* is the internal estimate of the motor state (i.e., the hand movement required to compensate for the perturbation), *a* is the retention factor, *e*(*n*) is the size of the perturbation in trial *n*, *b* is the error sensitivity for a given error size, and ε_x_ represents planning noise.

The actual motor response on trial *n* is given as:

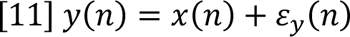

where *y* is the reaching direction relative to the target, determined by *x*(*n*) and motor noise, ε_-_.

For the dual state-space model, we added a second, slower learning process (*xs*) with a different retention rate (*as*) and learning rate (*bs*),

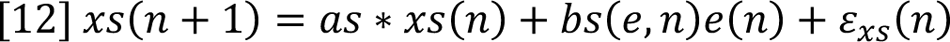

Where *as* > *a* and *bs* < *b*. As such, Eq [11] can be written as:

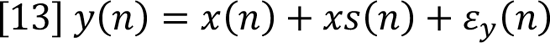

#### Memory-of-Error model (MoE)

The Memory-of-Error model describes how the learning rate in the state-space model is modulated by experience. In the MoE model, error sensitivity (*b*) is set to an initial value that is modulated by errors that are experienced during training. Specifically, *b(e,n)* will increase if the error on trial n+1 shares the same sign and *b(e, n)* will decrease if the error on trial n+1 is of the opposite sign. This is formalized as:

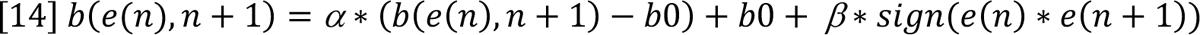

where *Β* and *α* are the learning rate and retention rate of *b*, respectively. Since the error size is fixed at 30° in our experiments, we replace *b*(*e*) with a single value *b*.

#### Contextual Inference (COIN) model

We simulated the Contextual Interference (COIN) using the code provided by Heald et al.^29^, focusing on its prediction with respect to savings and spontaneous recovery. We assumed that the introduction of a perturbation (e.g., clamped feedback) defines a new context and, as such, leads to the establishment of a new motor memory. Similarly, reversing the sign of the perturbation would define another context and thus require establishment of another memory. We simulated the clamps as if they were contingent rotations so that the learning can reach an asymptote. Before each movement, the output is determined by averaging the state of different contexts weighted by the expected probabilities of the contexts. Participants observed an error after each movement and update the state estimation.

### Behavioral Experiments

#### Participants

A total of 451 participants (297 female, mean age = 28.0, SD = 5.3) were recruited through the website prolific.co. After eliminating participants who failed to meet our performance criteria (2.8%, see below), the analyses were based on data from 438 participants. Based on a survey included in a prescreening questionnaire, the participants were right-handed with normal or corrected-to-normal vision. The participants were paid based on a rate of $8/h. The protocol was approved by the Institutional Review Board at the University of California, Berkeley. Informed consent was obtained from all participants.

#### Apparatus

All of the behavioral experiments were conducted online using a web-based experimental platform, OnPoint^58^, which is written in JavaScript and presented via Google Chrome. It is designed to operate on any laptop computer. Visual stimuli were presented on the laptop monitor and movements were produced on the computer trackpad. Data were collected and stored using Google Firebase.

#### Clamp rotation task

We applied clamp feedback in the experiments, under the assumption that learning in response to this type of feedback is limited to implicit, cerebellar-dependent sensorimotor recalibration. To start each trial, the participant moved the cursor to a white start circle (radius: 1% of the screen height) positioned in the center of the screen. After 500ms, the target, a blue circle (radius: 1% of the screen height) appeared with the radial distance set to 40% of the screen size. The target appears at −45°, a workspace location selected because it exhibits minimal bias across participants^80^. The participant was instructed to produce a rapid, out-and-back movement, attempting to intersect the target. If the movement time (from onset to time at which movement amplitude reached the target) was longer than 500ms, the message ‘Too Slow’ was presented on the screen for 500ms.

There were three types of feedback. On veridical feedback trials, the position of the cursor moved was matched to the position of the hand, subject to the translation in reference frames (screen assumed to be vertical, hand movement assumed to be horizontal) and scaling (trackpad space expanded to encompass most of the screen). On clamped feedback trials, the cursor followed a fixed path. As with veridical feedback, the radial location of the cursor was based on the radial extent of the participant’s hand. However, the angular position of the cursor was independent of the position of the hand, instead determined relative to the position of the target. The clamp angle was set at 30° relative to the target except for Exp 5 and 8 (see below). On no feedback trials, the cursor was blanked at movement onset.

On veridical and clamped feedback trials, after the amplitude of the movement reached the target distance, the cursor was presented at the target distance for another 50ms then it disappeared. Target disappeared after 200ms. The cursor was then reset to a random position on an invisible circle with a radius equal to 10% of the target distance and the participant moved the cursor back to the start circle.

At the onset of the first block of trials involving perturbed feedback, the experiment was paused, and a set of instructions were presented to describe the clamped feedback. The participant was informed that the cursor would no longer be linked to their movement but rather would follow a fixed path on all trials. The participant was instructed to always reach directly to the target, ignoring the cursor. These instructions were then repeated twice to emphasize the atypical nature of the feedback. After the first 10 trials with clamped feedback, a new instruction screen appeared in which the participant was asked to indicate where they were aiming on each trial. If the participant indicated they were reaching somewhere other than the target, the experiment was terminated.

Each experiment started with two baseline blocks: First a no-feedback block of 10 trials and second, a veridical feedback block of 10 trials. For experiments using a fixed design (direction and size of perturbation remain constant), the direction of the clamp (counterclockwise, CCW; clockwise; CW) was counterbalanced across participants.

#### Experiment 1

Exp 1 was designed to determine the parameters of the CPC model. There was a total of 180 trials. The two baseline blocks were followed by a learning block of 100 trials with clamped feedback with learning expected to reach an asymptotic level in response to a fixed perturbation. This was followed by a final no-feedback block of 60 trials. 30 participants were recruited for Exp 1 (29 valid, 5 males, age: 27.4 ± 4.9 years).

#### Experiment 2

Exp 2 was designed to measure antegrade interference. The baseline and initial perturbation blocks were as in Exp 1. For the final block (150 trials), the direction of the clamp was reversed (e.g., from 30° to −30°). 30 participants were recruited for Exp 6 (30 valid, 10 males, age: 30.3 ± 4.3 years).

#### Experiment 3

Exp 3 was designed to assess spontaneous recovery and savings in implicit adaptation. The baseline and initial perturbation blocks were as in Exp 2. We then included a 15-trial block with the clamp reversed under the assumption that this would be a sufficient number of trials to bring the hand angle back to baseline. This was followed by no-feedback block (35 trials) to examine spontaneous recovery and then a 100-trial relearning block in which the clamp feedback was identical to that used in the first perturbation block. 34 participants were recruited for Exp 3 (34 valid, 16 males, age: 22.7 ± 4.8 years).

#### Experiment 4

Exp 4 examined how the consistency of the perturbation influenced implicit adaptation. The first blocks were identical to Exp 3, providing initial exposure to clamped feedback and then a reversed clamp to bring the hand angle back to baseline. This was followed by a 300-trial block in which the clamp changed sign in a probabilistic manner. The probability of a sign change was either 90%, 50%, and 12.5% in a between-subject manipulation. The sequence of clamps was preset to ensure that clockwise and counterclockwise occurred on 50% of the trials each across the 300 trials. The experiment ended with a relearning block in which the initial perturbation was presented for 100 trials. 36/40/36 participants were recruited for 90%, 50%, and 12.5% conditions respectively (34/38/33 valid, 37 males, age: 28.6 ± 5.5 years).

#### Experiment 5

To estimate the learning rate and retention at top layer of the CPC model, the PF-PC synapse, we employed a variable design in which the error size and direction varied across trials. After the two baseline sections, participants completed a 540-trial random perturbation block. Here the clamp size ranged from −135° to 135° in steps of 1°. The size/direction was determined at random with the constraint that each clamp was selected once every 270 trials. 72 participants were recruited for Exp 5 (70 valid, 25 males, age: 26.2 ± 5.2 years).

#### Experiment 6

Exp 6 was designed to evaluate different models of asymptotic adaptation. A 10-trial feedback baseline block was followed by a learning block of 100 trials with clamped feedback. We then alternated between no-feedback and clamp feedback trials for 60 trials (half-wash phase). 40 participants were recruited for Exp 6 (38 valid, 8 males, age: 30.7 ± 6.6 years).

#### Experiment 7

Experiment 7 was designed to measure the time course of retention during the initial washout phase. After the two baseline blocks, the perturbation block consisted of 31 mini-blocks, each composed of 10 trials with clamped feedback and 10 trials without feedback (620 trials). 57 participants were recruited for Exp 7 (57 valid, 12 males, age: 28.3 ± 5.4 years).

#### Experiment 8

To quantify the temporal dynamics of volatile processes, we used variable clamped feedback with extended inter-trial intervals (ITI) in Exp 8. For the long ITI, the interval between the end of one trial and the start of the next trial was 6 s, 7 s, or 8 s, randomized across trials. The message “wait” was displayed on the monitor after each trial. Exp 8 included two baseline blocks and a 180-trial learning block in which a 30° perturbation was randomly selected to be either clockwise or counterclockwise, subject to the constraint that each direction occurred four times every 8 trials. For the short ITI condition, we used the data from Exp 4 for the variable condition (0 s ITI). 28 participants were recruited for each condition (27 valid, 13 males, age: 28.1 ± 4.8 years).

#### Experiment 9

To understand how the volatile and stable learning processes are jointly modulated by time, we used a fixed design in Exp 9. The design was similar to that employed in Exp 1 with one notable modification. We included a 10-trial familiarization block following the two baseline blocks to demonstrate the clamp feedback. The clamp size in the familiarization block varied from −90° to 90° across trials to show that the cursor is unaffected by the direction of hand movement. To avoid the influence of pre-exposure to the error signal on learning, the familiarization block utilized a different target (45°) from the other blocks (315°). Two groups of participants performed the task with either long ITI (6-8s) or short ITI (0s). 26 participants were recruited for each condition (51 valid, 21 males, age: 26.8 ± 4.6 years).

### Data analyses

Hand angle was calculated as the angle difference a line from the start position to the target and a line from the start position to the hand position at the target radius. Positive values indicate hand angles in the opposite direction of the perturbation, the direction one would expect due to adaptation. Trials with a movement duration longer than 500 ms or an error larger than 70° were excluded from the analyses.

We excluded the entire data from participants who had less than 70% valid trials (2.8% participants). Between-condition comparisons were performed with t-tests or ANOVAs. Learning and relearning were compared by paired-t-test. For all the statistical tests, we confirmed that the data met the assumptions of a Gaussian distribution and homoscedasticity.

### Data and Software Availability

The data and code supporting this work are available at https://github.com/shion707/CPC.

## Author contributions

T.W., R.B.I. contributed to the conceptual development of this project. T.W. collected the data, analyzed the data, prepared the figures, and wrote the initial draft of the paper, with both authors participated in the editing process.

## Acknowledgment

This study is funded by the NIH (grants NS116883 and NS105839). We thank Chris Miall for helpful discussion.

## Supplementary Information

### Supplemental Result 1: The CPC model accounts for contextual interference

The two-layer model provides an alternative explanation for another type of context-dependent learning, contextual interference. The term is a bit of a misnomer since the phenomenon refers to the fact that, while performance gains when training in multiple contexts is slower compared to training in a single context, retention is better in the former^69,70^. As such, exposure to multiple contexts during training actually enhances learning as measured by long-term gains. Interestingly, this phenomenon is not limited to skill acquisition tasks but is also observed in studies of implicit adaptation^81^ (Fig S9).

In the revised CPC model, contextual interference occurs due to the parallel operation of volatile and stable learning processes. With multiple targets (constituting multiple contexts), the rate of acquisition is slower compared to a single target since learning from the volatile process decays between successive reaches to a given target. However, early retention is higher since the contribution of the volatile process is small. Thus, as with anterograde interference, contextual interference arises from the dynamics of the system without postulating any representation of context.

---

**Fig. S1.**
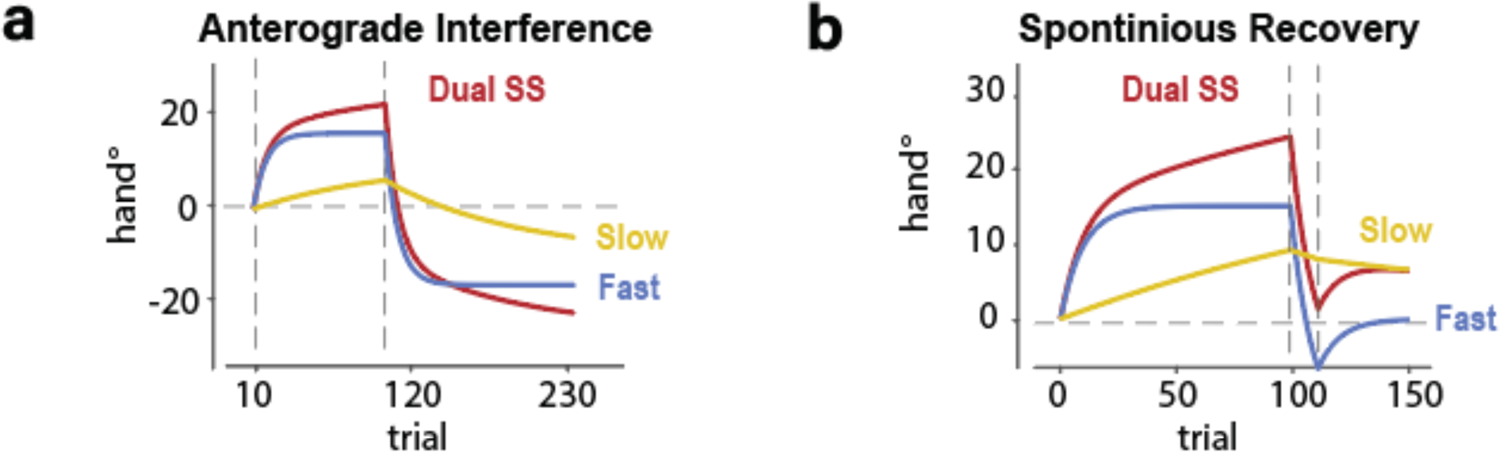
Predictions of the Dual State Space model for Exp 2-3. The time course of the predicted hand angle as well as the underlying states of the fast and slow processes are shown.

**Fig. S2.**
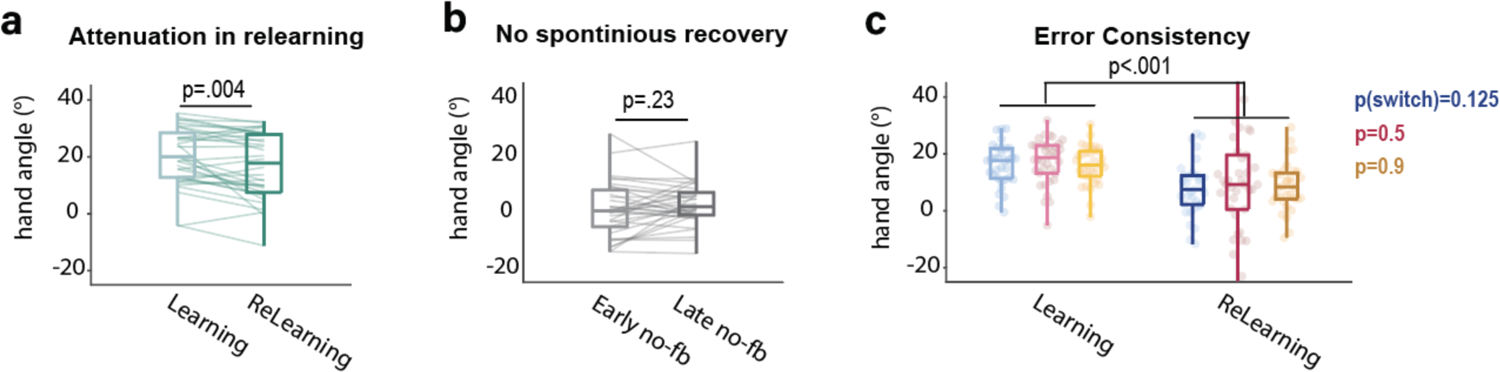
Effect of experience and error consistency on implicit adaptation. **a)** Attenuation in relearning in Exp 3. Adaptation was attenuated in response to re-exposure to a perturbation compared to the initial exposure (t(33)=3.1, p=0.004) Data are averaged across each training phase. **b)** Spontaneous recovery was not observed in Exp 3 during the no-feedback phase after washout. Hand angle over the first 5 trials of the no-feedback phase (Early) is similar to hand angle over the last 5 trials (Late, t(33)=1.2, p=0.23). c) Error consistency did not affect adaptation during initial learning and during relearning in Exp 4. A mixed ANOVA showed a main effect of learning/relearning, (F(1,101)=37.7, p<0.001), similar to the antegrade interference observed in Exp 6. There was no effect of error consistency (F(2,101)=0.18, p=0.84) or interaction between phase and error consistency (F(2,101)=0.12, p=0.88). Box plots indicate median, max and min values, and 25% and 75% quartiles.

**Fig. S3.**
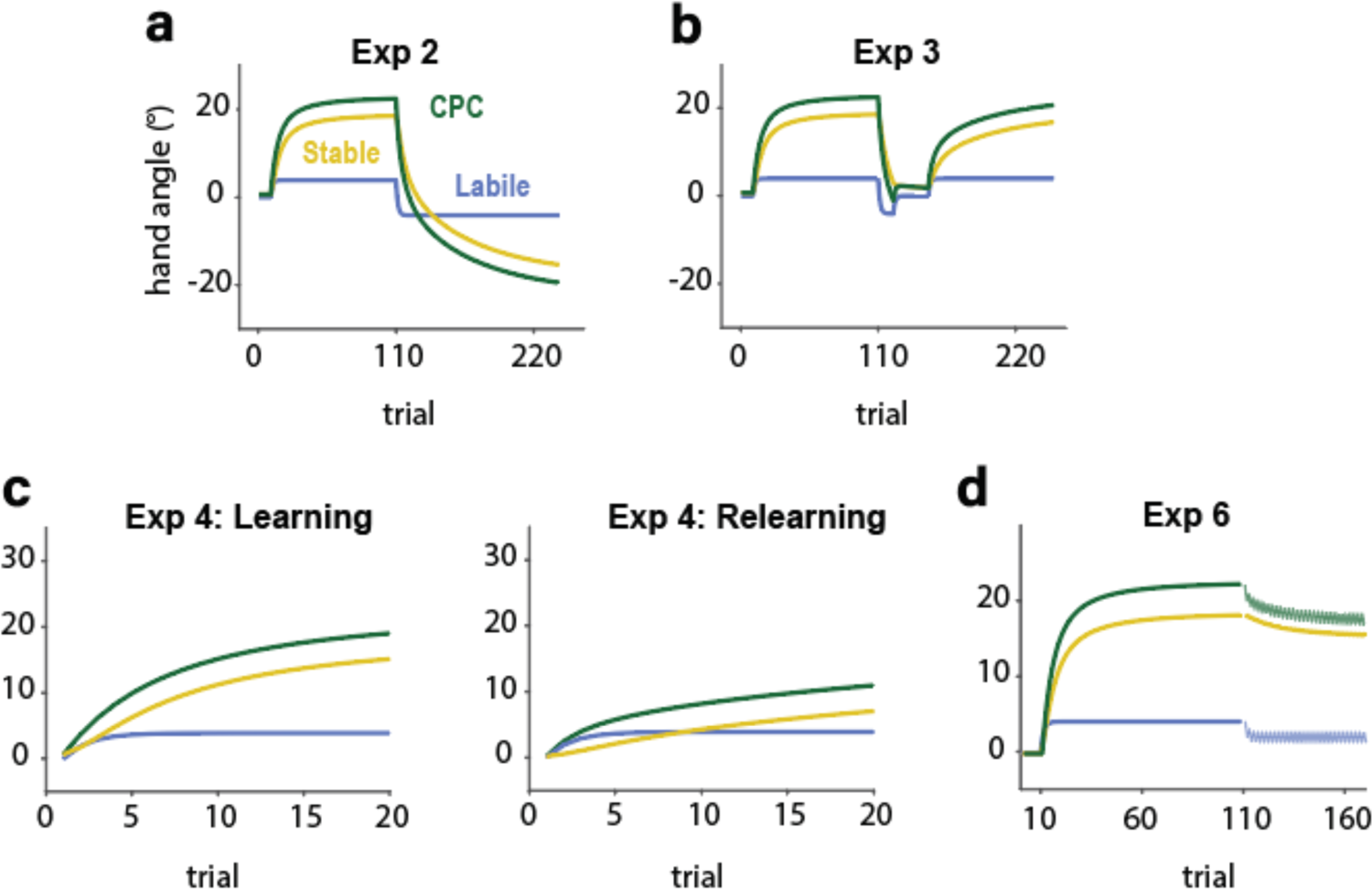
Predicted time course of stable and volatile processes in Exps 2-4 and 6. The stable process is responsible for anterograde interference (a) and attenuation in relearning (b-c). The volatile process does not make a significant contribution to either phenomenon because of its low retention rate.

**Fig. S4.**
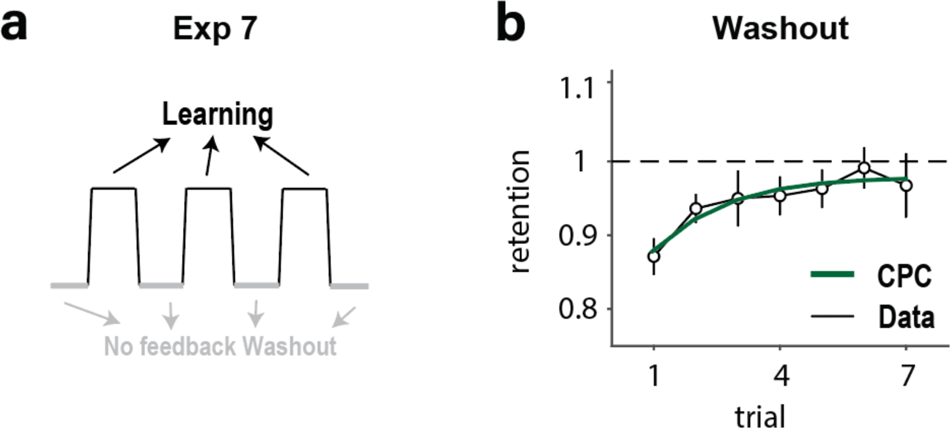
Retention increases during the initial washout trials. **a)** To provide a stronger test of how the rate of retention changes (Exp 1), Exp 7 included mini-blocks (10 trials/mini-block) that alternated between clamp and no feedback trials. **B**) We estimated the change in retention rate over time by averaging by trial number across the no feedback blocks. Retention is relatively low in the first trials of the washout block and gradually rises (F(6,264)=4.64, p<0.001). The dark green curve shows the fit of the CPC model.

**Fig. S5.**
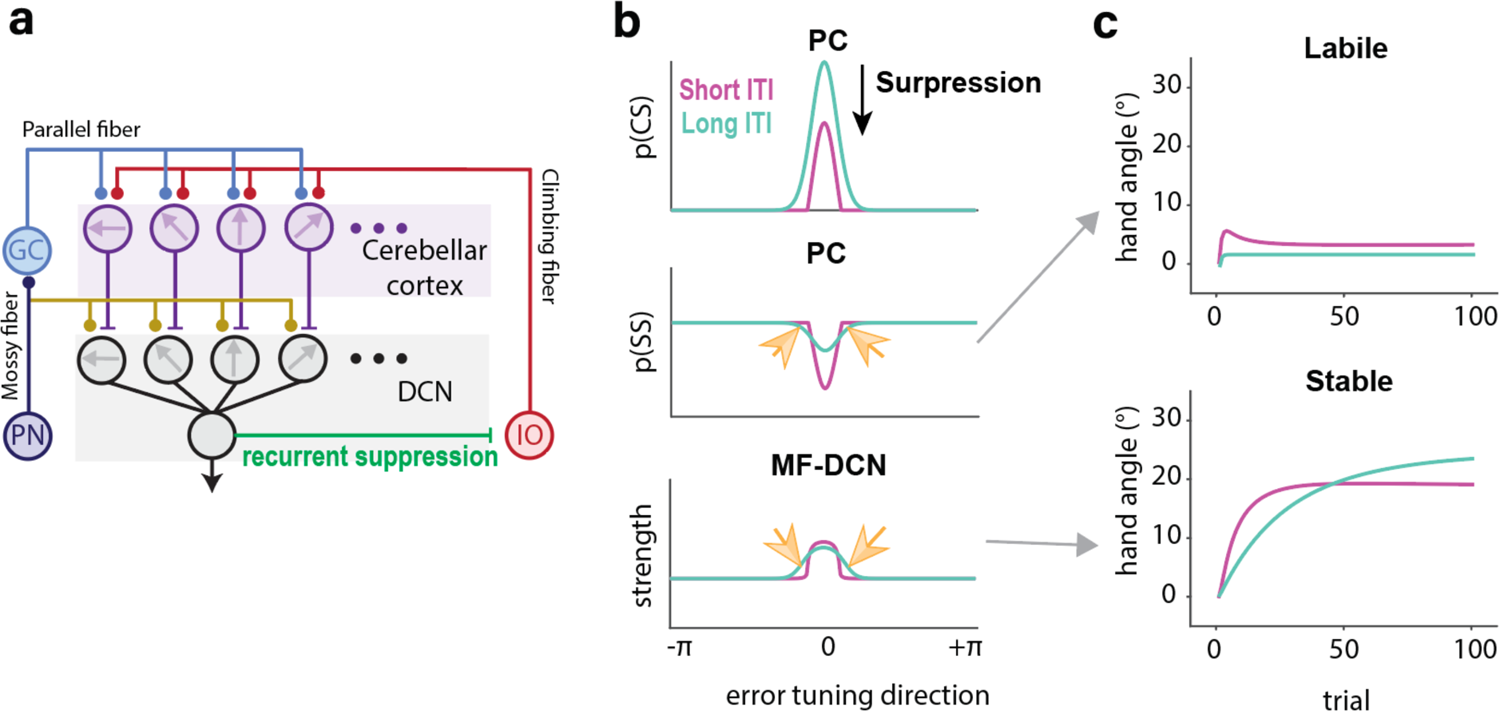
Revised CPC with DCN-IO inhibitory pathway. The original CPC predicts that the asymptote should be lower in the long ITI condition compared to the short ITI condition because the latter includes a volatile component. However, as shown in Fig 7, the asymptote is similar in the two ITI conditions. This observation motivated a revision to the CPC model in which the DCN sends a inhibitory signal to the inferior olive. **a)** Model schematic. DCN-IO inhibition suppresses the error signal to the DCN and cerebellar cortex. This suppression is generic given that the output of the DCN integrates activation across directionally tuned units. **b)** When the inter-trial-interval is short, the CS response is suppressed (top). Note that the suppression is implemented by subtracting a common value to the IO and thus alters the activation in PCs. On the next trial, SS activation is stronger in the long ITI condition since the PF-PC synapse will have recovered during the ITI (middle). However, there are a subset of tuned elements that in which SS activation is weaker in the long ITI condition (yellow arrows). This weaker activation induces adaptation in DCN units tuned to the same direction (bottom). **c)** State of the volatile and stable processes over the course of a fixed design under long and short ITI conditions. The change in the volatile process is smaller in the long ITI condition due to forgetting. The stable process is also smaller in the long ITI condition because SS activity at the preferred error direction will dominate learning. However, the long ITI condition induces adaptation in neurons with sub-preferred error directions, resulting in larger adaptation late in training.

**Fig. S6.**
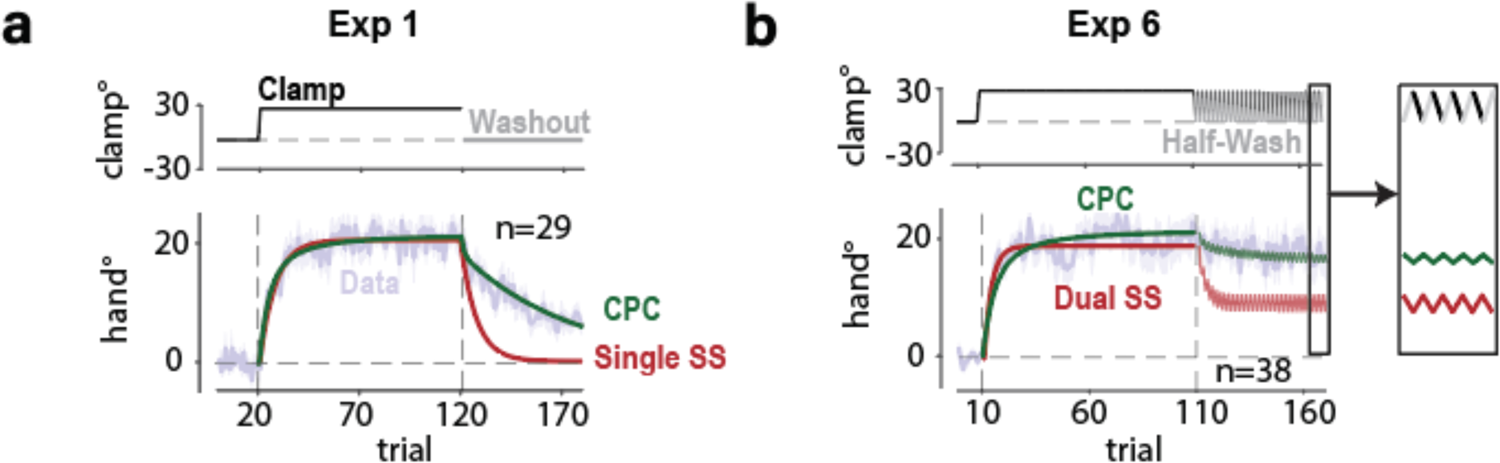
State-space models fails to explain the learning asymptote. **a)** A single state motor cannot account for fast early learning and slow forgetting. **b)** The state-space model assumes the asymptote reflect a balance between learning and forgetting. As such the asymptote will drop to a half in the half washout phase of Exp 6. However, there is only a slight decrease in the asymptote during the half washout, consistent with the predictions of the CPC model rather than the dual SS model.

**Fig. S7.**
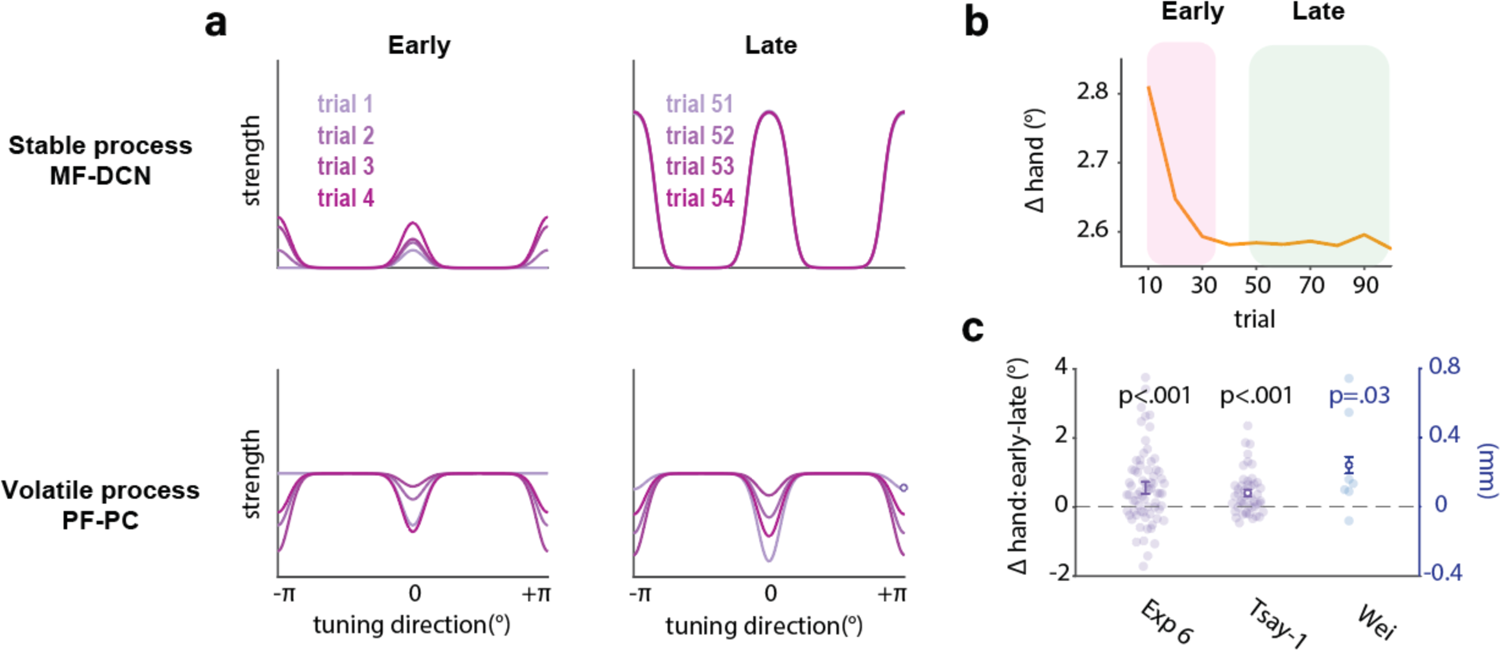
Contribution of stable and volatile processes in response to variable perturbations. **a)** The stable process (top) contributes to learning during early training and has saturated by the 50^th^ trial. The contribution of the volatile process (bottom) remains similar throughout training. **b)** Change in hand angle as a function of trial number when the size and direction of the perturbation varies across trials. The change of hand angle is larger in early training because the stable process has not saturated. **i)** As predicted by the two-process CPC model, when exposed to a variable perturbation, the Δhand is larger in early training compared to late training. Shaded areas and error bars indicate standard error.

**Fig. S8.**
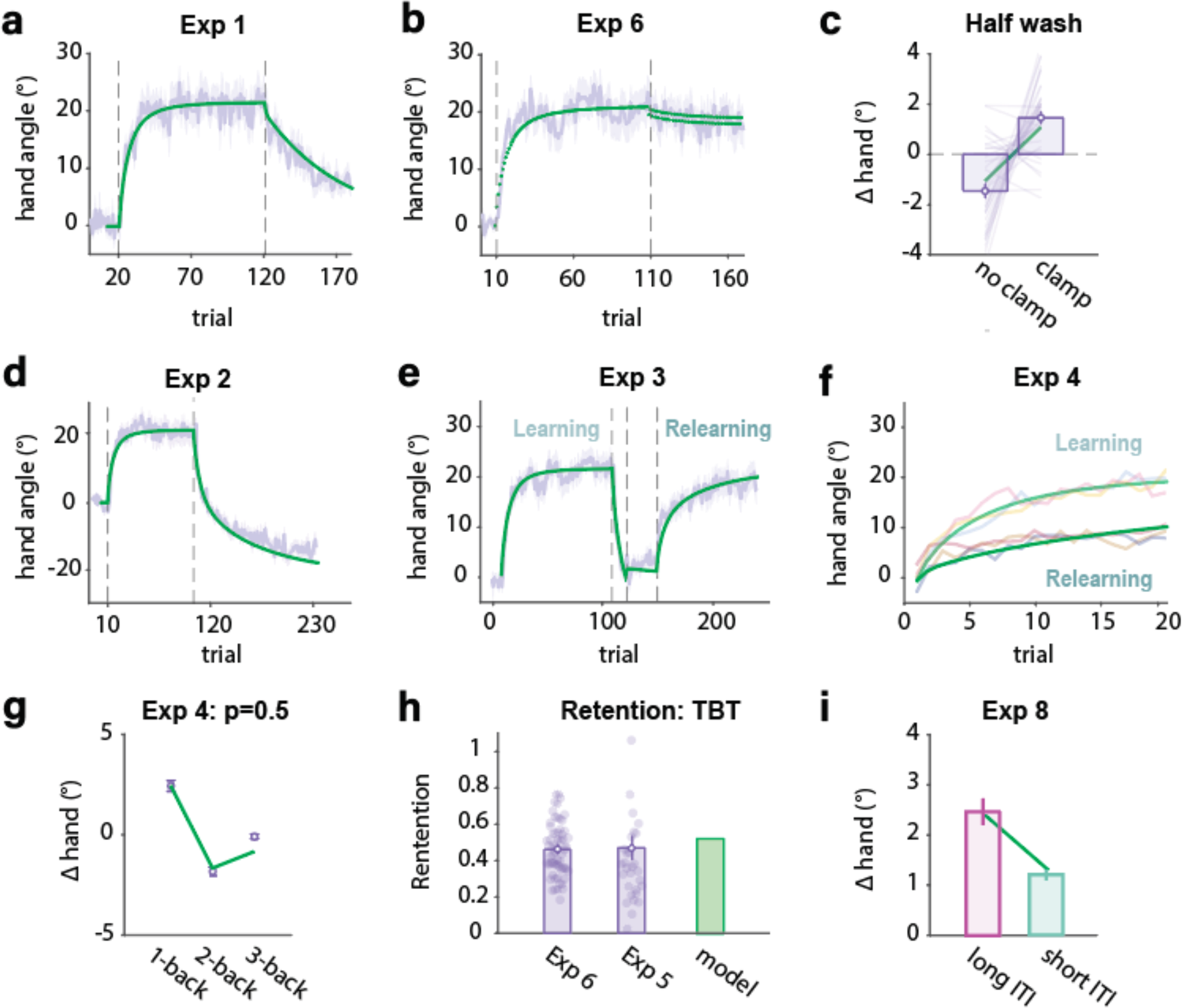
Revised CPC model provides a good fit for the key results for all of the experiments. Dark green line depicts model prediction. Error bars (c, g, h, i) and shaded areas (a, b, d, e) indicate standard error.

**Fig. S9.**
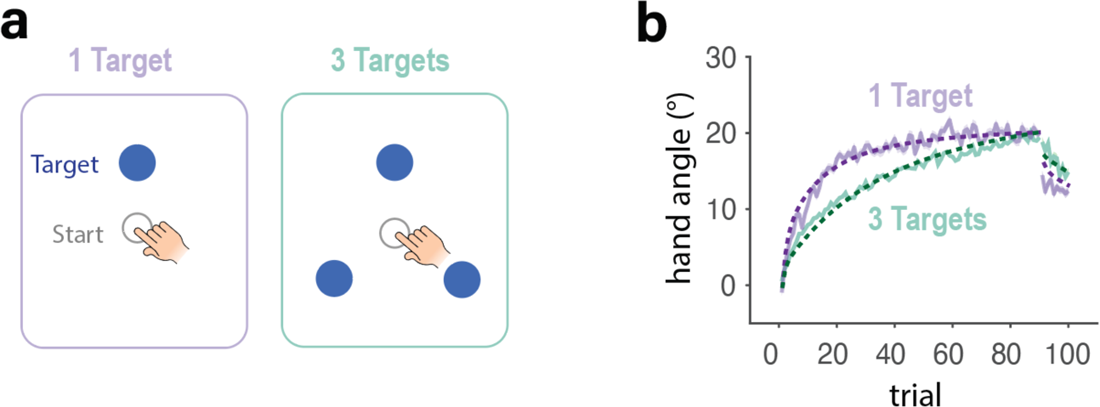
Revised CPC model accounts for effect of number of target locations on adaptation. **a)** In Tsay et al.^82^, participants were trained with either one target or three targets. In both conditions, participants reached to a single target during the washout block. **b)** Learning functions for the target location probed during washout. The 3-target condition showed slower learning but a larger aftereffect. Adding more targets is effectively akin to imposing a long ITI since successive reaches to a given target are separated by reaches to the other two locations; thus, there is more forgetting but stronger retention due to reduced contribution of volatile process. Shaded area in b indicates standard error. Dash lines indicate the predictions of the Revised CPC model.

